# Quantitative yeast-yeast two hybrid for discovery and binding affinity estimation of protein-protein interactions

**DOI:** 10.1101/2020.08.12.247874

**Authors:** Kaitlyn Bacon, Abigail Blain, John Bowen, Matthew Burroughs, Nikki McArthur, Stefano Menegatti, Balaji M. Rao

## Abstract

Quantifying the binding affinity of protein-protein interactions is important for elucidating connections within biochemical signaling pathways, as well as characterization of binding proteins isolated from combinatorial libraries. We describe a quantitative yeast-yeast two hybrid (qYY2H) system that not only enables discovery of specific protein-protein interactions, but also efficient, quantitative estimation of their binding affinities (*K*_*D*_). In qYY2H, the bait and prey proteins are expressed as yeast cell surface fusions using yeast surface display. We developed a semi-empirical framework for estimating the *K*_*D*_ of monovalent bait-prey interactions, using measurements of the apparent *K*_*D*_ of yeast-yeast binding, which is mediated by multivalent interactions between yeast-displayed bait and prey. Using qYY2H, we identified interaction partners of SMAD3 and the tandem WW domains of YAP from a cDNA library and characterized their binding affinities. Finally, we showed that qYY2H could also quantitatively evaluate binding interactions mediated by post-translational modifications on the bait protein.

## Introduction

Quantitative binding affinity estimation of protein-protein interactions (PPIs) is important for system-level analysis of intracellular biochemical signaling pathways (1–6), as well as characterization of binding proteins isolated from combinatorial libraries using screening platforms such as phage display (7, 8) and yeast surface display (9, 10). Yeast two hybrid (Y2H) assays are a powerful means to discover putative protein interaction partners (“prey”) for specific proteins of interest (“bait”) in a high throughput manner (11, 12). Modifications to Y2H have enabled the elucidation of interactions that rely on post-translational modifications by tethering a modifying enzyme to the bait protein (13, 14), as well as semi-quantitative characterization of relative binding affinities using yeast surface displayed bait proteins (15). More recently, yeast cell surface expression of bait and prey proteins as fusions to yeast mating proteins has also been used to identify PPIs in a high throughput fashion; the quantitative relationship between binding affinities and efficiencies of yeast mating can be used to assess the strength of bait-prey binding (16). However, despite these advances, obtaining quantitative estimates of binding affinity – specifically equilibrium dissociation constant (*K*_*D*_) values – is generally not feasible using the aforementioned approaches.

Here, we describe a quantitative yeast-yeast two hybrid (qYY2H) system that enables the discovery of protein interaction pairs, as well as efficient and quantitative estimation of interaction binding affinities (*K*_*D*_). The requirement for recombinant, soluble protein is often a significant hurdle for high throughput binding affinity estimation (17, 18). In qYY2H, both the bait and prey proteins are expressed as cell surface fusions using the widely implemented Aga2-based yeast display platform (9, 10), which bypasses the need for recombinant protein **(Fig. 1)**. Bait cells co-express the iron oxide binding protein, SsoFe2, as a cell surface fusion, enabling magnetization of bait yeast and separation of bait-prey complexes (19, 20). An engineered luciferase reporter, NanoLuc (21, 22), is expressed on the surface of prey yeast that can be used to quantify the number of prey cells complexed with the bait cells, and therefore, the apparent *K*_*D*_ describing yeast-yeast binding interactions that are mediated by multivalent associations between bait and prey proteins expressed on each yeast population. Importantly, we describe a semi-empirical framework that enables quantitative estimation of the *K*_*D*_ of monovalent interactions between the bait and prey proteins using estimates of multivalent yeast-yeast binding affinities. Further, we show that qYY2H can be used to efficiently identify and quantitatively characterize putative interaction partners of bait proteins from a yeast displayed cDNA library. Finally, quantitative assessment of binding affinities is particularly challenging using conventional methods when one or both proteins under investigation are post-translationally modified. We have modified qYY2H to address this challenge by expressing and sequestering the modifying enzyme in the endoplasmic reticulum (ER), which can act on the expressed bait protein prior to surface display. Using this approach, we show that qYY2H can be used to quantitatively characterize binding interactions mediated by phosphorylated bait.

**Figure 1.**
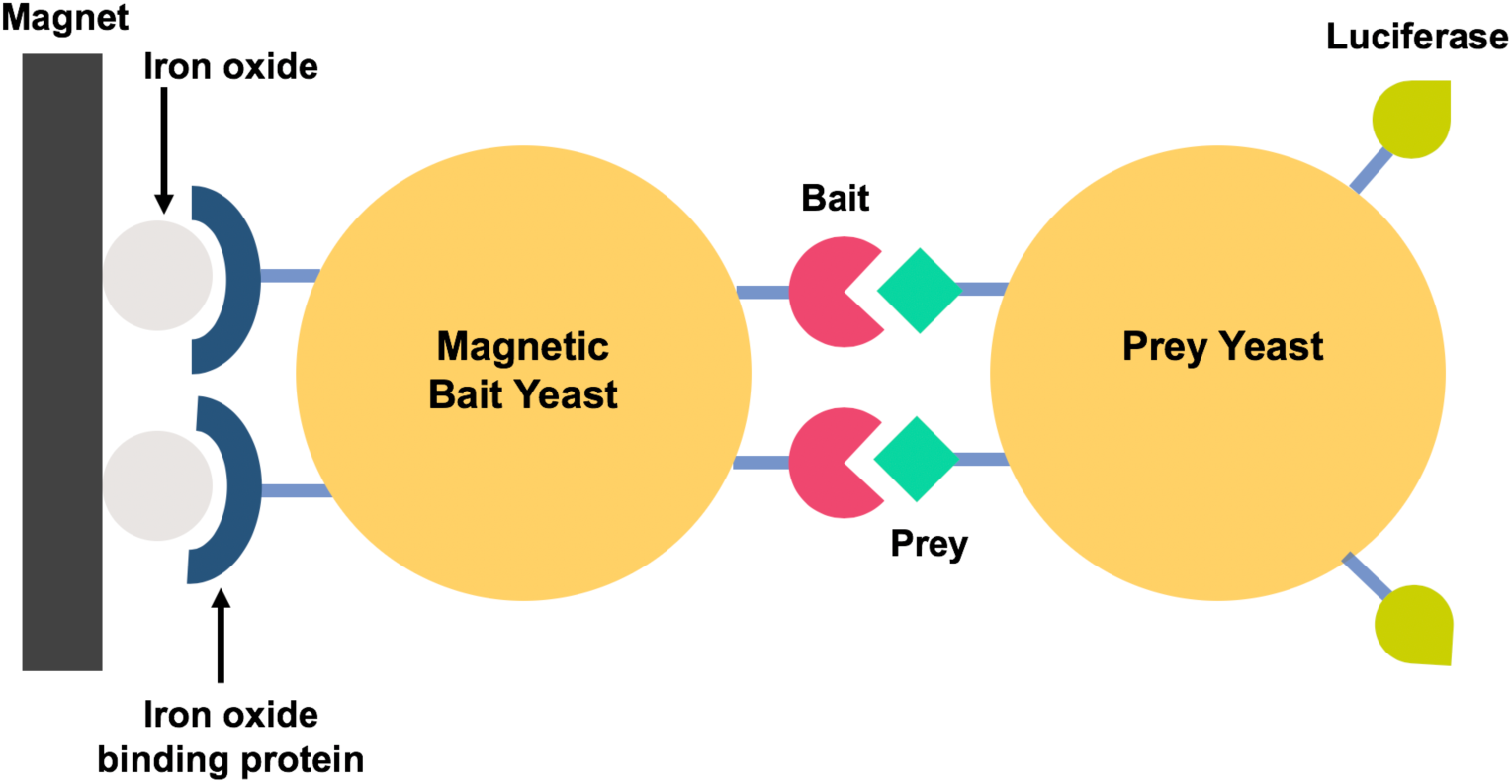
Quantitative yeast-yeast two hybrid (qYY2H). Yeast cells co-expressing the bait protein and SsoFe2 are magnetized by incubation with iron oxide particles. Magnetic separation is used to isolate yeast cells expressing prey proteins that interact with the bait cells. Prey yeast also display a luciferase reporter. The number of prey cells captured can be accurately quantified using a luminescence assay, in conjunction with a standard curve.

## Results and Discussion

### qYY2H enables relative binding affinity discrimination between bait-prey interactions

In qYY2H, the interaction of magnetized bait yeast with prey cells results in the formation of bait-prey, yeast-yeast complexes that can be separated using a magnet **(Fig. S1)**. Subsequently, the number of prey cells captured can be accurately quantified using a luminescence assay. We hypothesized that the number of prey cells captured can be used to quantitatively discriminate between the relative strengths of bait-prey binding interactions. To test this hypothesis, we investigated, in the context of qYY2H, the interaction of two previously characterized binding proteins (prey) based on the Sso7d protein scaffold – Sso7d.hFc (*K*_*D*_ = 450 nM) and Sso7d.ev.hFc (*K*_*D*_ = 5280 nM) – with the Fc portion of human immunoglobulin G (hFc; bait). The number of Sso7d.hFc prey cells captured was ∼3-fold higher than the number of Sso7d.ev.hFc prey cells recovered **(Fig. 2A)**. Further, the number of cells captured displaying the lower affinity Sso7d.ev.hFc was ∼ 4-fold higher than background (number of cells displaying an irrelevant prey protein). These results show that qYY2H can be used to discriminate between the relative strengths of bait-prey interactions. It is important to note that the high sensitivity of the NanoLuc reporter enzyme is critical for discrimination of relative binding strengths in qYY2H. Relative binding strengths of bait-prey interactions could not be assessed when glucose oxidase (GOx) was used as a reporter **(Fig. S2)**. Additionally, non-specific binding between yeast cells is common. Therefore, inclusion of excess competitor, non-displaying EBY100 yeast and a non-ionic detergent, such as Tween-20, is important **(Fig. S1)**.

**Figure 2.**
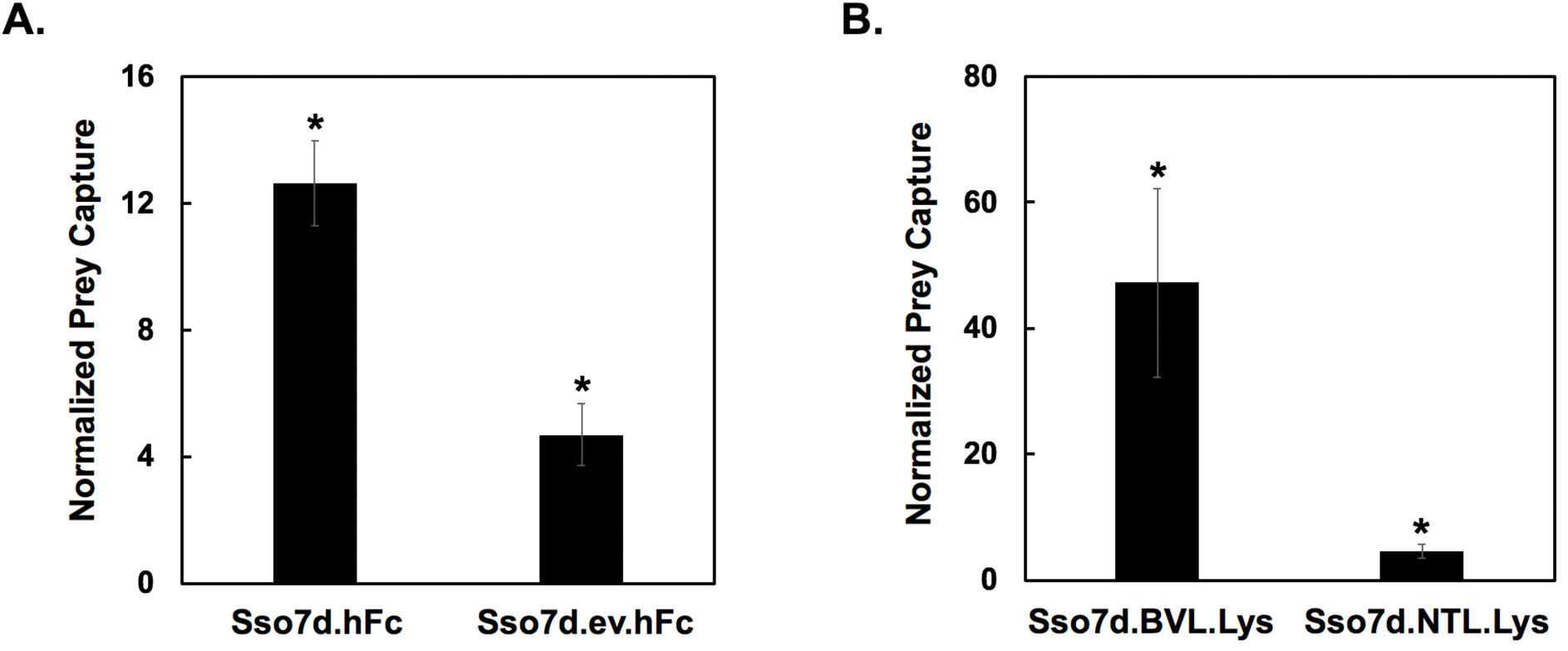
qYY2H enables discrimination of relative binding affinities associated with bait-prey interactions through display of a NanoLuc reporter. **(A)** Recovery of prey cells by magnetic yeast displaying the bait protein (hFc). The prey cells displayed either Sso7d.hFc, Sso7d.ev.hFc, or an irrelevant protein, NanoLuc. Recovery of the cells displaying the hFc binding proteins was normalized by the capture of cells displaying NanoLuc. * represents p < 0.05 for a two tailed, paired t-test in comparison to the capture of NanoLuc cells. **(B)** Recovery of prey cells by magnetic beads immobilized with the bait protein (lysozyme). The prey cells displayed either Sso7d.BVL.Lys, Sso7d.NTL.Lys, or an irrelevant protein, Sso7d.hFc. Recovery of the cells displaying the lysozyme binding proteins was normalized by the capture of cells displaying Sso7d.hFc. *represents p < 0.05 for a two tailed, paired t-test in comparison to the capture of Sso7d.hFc cells. Error bars represent the standard error of the mean for three replicates.

Interestingly, quantitative discrimination of relative binding strengths was also observed when yeast cells displaying the lysozyme-binding proteins Sso7d.BVL.Lys (*K*_*D*_ = 1.7 nM) and Sso7d.NTL.Lys (*K*_*D*_ = 1300 nM) were used as prey, and biotinylated bait protein (lysozyme) was immobilized on streptavidin coated magnetic beads **(Fig. 2B)**. Recovery after magnetic separation was ∼ 20-fold greater for cells displaying the higher affinity Sso7d.BVL.Lys. Our results are strikingly different from those previously reported by Ackerman *et al* (23). In that study, the recovery of cells displaying lysozyme-binding proteins was not affected by the displayed protein’s binding affinity for lysozyme, which was similarly immobilized on streptavidin-coated magnetic beads. A likely explanation for this discrepancy is the lower ratio of cells to magnetic beads used in our study. Taken together, our results show that the efficiency of magnetically isolating prey cells – using magnetized yeast cells displaying the bait or magnetic beads coated with bait – has a quantitative dependence on the binding affinity of the bait-prey interaction. Notably, this observation is contrary to the generally accepted view that magnetic selections, which are employed during combinatorial screening of yeast display libraries, do not discriminate between high and low affinity binders.

### A semi-empirical framework enables estimation of K_D_ using qYY2H

Formation of bait-prey, yeast-yeast complexes in qYY2H is driven by high avidity interactions between surface displayed bait and prey proteins. The number of bait-prey, yeast-yeast complexes isolated by magnetic separation was related to the apparent affinity of multivalent interaction between bait and prey yeast (*K*_*D,MV*_) using a monovalent binding isotherm, previously applied when performing yeast surface titrations with soluble prey protein (9):

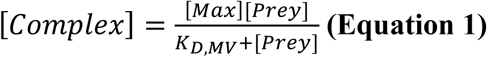

Where [*Complex*] is the molar concentration of bait-prey, yeast-yeast complexes, [*Prey*] is the initial molar concentration of prey yeast used in the experiment, and [*Max*] is the maximum molar concentration of the prey cells captured for a given number of magnetized bait yeast. We generated titration curves for seven bait-prey pairs where varying concentrations of prey cells (10^5^ – 4×10^8^ cells) were incubated with 10^7^ magnetized bait cells in the presence of 10^9^ non-displaying EBY100 yeast. The bait-prey pairs comprise binders to hFc and TOM22 based on the Sso7d or nanobody scaffolds and span a range of binding affinities (*K*_*D*_ ∼ 200 nM to *K*_*D*_ ∼ 5 μM). Binding affinities were previously estimated via yeast surface titration with soluble protein. *K*_*D,MV*_ and [*Max*] were estimated for each bait-prey pair **(Figs. 3A-E, S3)** by non-linear regression using **Equation 1**. It is important to note that saturation was not observed in several titration curves, particularly for higher affinity bait-prey pairs; increasing the concentration of incubated prey yeast further to saturate the binding of the bait yeast is not feasible in these cases.

**Figure 3.**
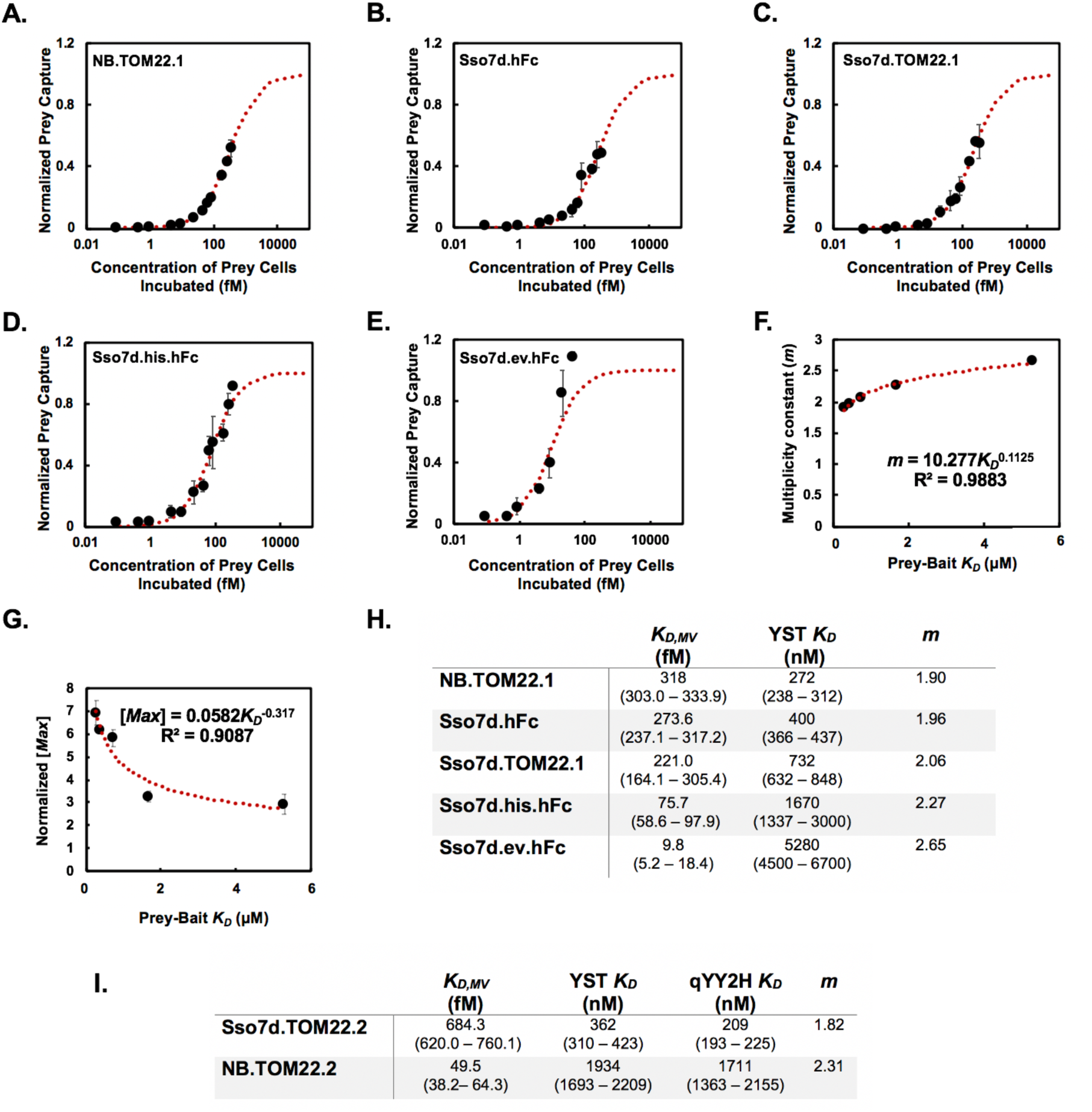
A semi-empirical framework enables estimation of the equilibrium dissociation constant of monovalent bait-prey binding (*K*_*D*_) using qYY2H. Titration curves describing the binding of prey cells as a function of the concentration of prey cells incubated were generated for a given concentration of magnetic bait cells. For each repeat, the concentration of prey cells captured is normalized by the estimated maximum concentration of prey cells captured, [*Max*]. Titration curves are shown for **(A)** NB.TOM22.1, **(B)** Sso7d.hFc, **(C)** Sso7d.TOM22.1, **(D)** Sso7d.his.hFc, and **(E)** Sso7d.ev.hFc. **(F)** A global, non-linear regression was used to estimate *K*_*D,MV*_, the equilibrium dissociation constant describing bait-prey, yeast-yeast binding. Associated multiplicity constants, *m*, were calculated using the fitted *K*_*D,MV*_ values and experimental *K*_*D*_ values. A plot of *m* vs. *K*_*D*_ is shown and fits a power law model. *K*_*D*_ values for these bait-prey interactions were previously estimated from soluble yeast surface titration (YST *K*_*D*_). **(G)** A plot of normalized [*Max*] values vs. *K*_*D*_ fits a power law model. [*Max*] values for each repeat were normalized by the average number of prey cells captured across each concentration of prey cells incubated. **(H)** Fitted values of *K*_*D,MV*_ and calculated values of *m* based on previous estimates of *K*_*D*_ (YST *K*_*D*_) are shown. **(I)** The *K*_*D*_ of Sso7d.TOM22.2 and NB.TOM22.2 is 209 nM (193 – 225 nM, 68% confidence interval) and 1711 nM (1363 – 2155 nM, respectively, as estimated using the multivalent binding model (qYY2H *K*_*D*_), compared to previously estimated *K*_*D*_ values of 362 nM (310 – 423 nM) and 1934 nM (1693 – 2209 nM) using soluble yeast surface titrations (YST *K*_*D*_). Values in parentheses represent the bounds of a 68% confidence interval. Error bars represent the standard error of the mean for three repeats for all data shown.

We hypothesized that a quantitative relationship exists between the binding affinity of the monovalent bait-prey interaction (*K*_*D*_) and the apparent affinity of the multivalent interaction between bait and prey yeast (*K*_*D,MV*_), similar to a previously outlined relationship that describes the multivalent interaction between viral proteins and cell surface receptors (24):

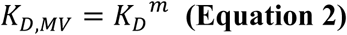

where *m* is a multiplicity constant that captures the increase in apparent affinity due to multivalent interaction. We used fitted values of *K*_*D,MV*_ and previously estimated values of *K*_*D*_ to calculate *m* for five randomly chosen bait-prey pairs **(Fig. 3H)**. We observed that *m* differed with *K*_*D*_, and this variation could be described using a power law relationship **(Fig. 3F)**, with *m* monotonically increasing with *K*_*D*_. Thus, effective increases in affinity due to multivalency is greater for bait-prey interactions with lower binding affinity. We also observed a power law relationship between [*Max*] and previously estimated values of *K*_*D*_ **(Fig. 3G)**. Lower affinity prey exhibited lower [*Max*] values than higher affinity prey. This may be explained by greater losses of bait-bound prey cells after the wash steps, due to weaker bait-prey, yeast-yeast interaction. Notably, **Equation 2** and the relationship developed between *m* and *K*_*D*_ **(Fig. 3F)** can be used to estimate *K*_*D*_ values using experimentally determined values of *K*_*D,MV*_ obtained from yeast-yeast titrations. Using this approach, *K*_*D*_ values were calculated for two binder-prey interactions randomly designated as “unknowns”. The obtained *K*_*D*_ values were in reasonable agreement with *K*_*D*_ estimates from yeast surface titrations using soluble prey protein **(Fig. 3I)**. These results show that qYY2H can be used to obtain quantitative estimates of binding affinity (*K*_*D*_) using yeast-displayed bait and prey proteins.

The strength of the multivalent interaction between bait and prey yeast will likely depend on the cell surface density of bait and prey proteins. In our studies, a robust correlation between *m* and *K*_*D*_ was obtained **(Fig. 3F)** even though the expression level of bait proteins on the yeast cell surface varied ∼3 fold across our studies **(Fig. S4)**. Therefore, the relationship between *m* and *K*_*D*_ described in **Fig. 3F** may be used to obtain reasonable estimates of *K*_*D*_ for other bait-prey interactions, despite some variation in cell surface display levels. To further investigate the effect of bait surface density, we conducted titrations with three hFc binding prey proteins displayed on the yeast surface using hFc, immobilized on magnetic beads, as bait **(Fig. S5)**. The surface density of hFc is significantly higher on magnetic beads (>2×10^5^ based on the manufacturer’s capacity for IgG binding) compared to level of hFc expressed on the yeast surface (∼5×10^4^) (9). We observed that *K*_*D,MV*_ is significantly lower for yeast-bead interactions relative to yeast-yeast interactions **(Fig. S6)**, indicating that the multiplicity constant *m* is greater when the surface density of the bait protein is higher. This is consistent with previous studies which show that *m* for virus-cell interactions is higher for cases where the expression level of virus-binding receptors on the cell surface is greater (24).

### qYY2H enables screening of cDNA libraries to identify putative PPIs and estimation of their binding affinities

Y2H is routinely utilized to identify putative interaction partners for a given bait protein by employing a cDNA-encoded protein library (25–27). To assess if qYY2H can be used for discovery of PPIs, we screened a cDNA library to identify proteins that interact with SMAD3, a signaling molecule that transduces TGF-beta signals from the cell surface to the nucleus by acting as a transcriptional regulator for various genes (28). Briefly, we constructed a yeast displayed cDNA library that expresses prey proteins as well as NanoLuc as cell surface fusions. The bait protein, SMAD3, was co-expressed with the iron oxide binding protein SsoFe2 to enable magnetization of bait yeast. Selections were conducted by incubating the yeast cells displaying prey proteins from the cDNA library with magnetized bait cells, followed by magnetic separation of bait-prey complexes, and selective expansion of positively bound prey yeast. After two rounds of selection, plasmid DNA was isolated from individual prey yeast clones. DNA sequencing of thirty clones identified nine unique sequences corresponding to putative SMAD3-binding proteins **(Table S1)**. Note that we also recovered prey cells displaying a Sso7d mutant protein that arises from incomplete digestion of the plasmid vector used as the starting backbone to construct the cDNA library. A likely explanation for recovery of these cells is association between the Sso7d prey mutant and the SsoFe2 protein used for magnetizing bait cells. Importantly, the inadvertently included Sso7d-displaying prey cells act as competitor for the cDNA clones, thereby increasing the likelihood that the putative cDNA prey isolated do not bind non-specifically to the bait protein.

We compared the interaction strength between SMAD3 and a subset of the identified putative binding partners, with that of the SMAD3-SARA binding interaction. SARA binds unphosphorylated SMAD3 (29–31) and thus acts as a positive control. Specifically, we quantified the number of prey cells captured by magnetized yeast cells displaying SMAD3, relative to the number of cells captured displaying the SMAD binding domain of SARA **(Fig. 4A)**. Cells displaying a portion of the cohesin loading factor NIPBL protein were captured at the highest rate by the SMAD3 bait cells, followed by cells displaying fragments of bromodomain containing protein 7 (BRD7), gametogenetin binding protein 2 (GGNBP2), and endothelin-1 (END1). In comparison to cells displaying SARA, cells expressing each of these prey were captured at a higher level, suggesting that the binding strength of these protein fragments for SMAD3 is at least comparable to that of SARA. On the other hand, fewer cells displaying portions of the mitochondrial calcium uniporter (MCU) protein and the dead box polypeptide 18 (DDX18) protein were recovered than cells displaying SARA, suggesting that these proteins may bind SMAD3 with lower affinity than SARA. Notably, BRD7 has been previously shown to interact with SMAD3 (32). Similarly, other RNA helicase dead box proteins, similar to DDX18, have been shown to complex with SMAD3 (29, 33). However, two of the identified putative binding partners – MCU and endothelin 1 – are localized to the mitochondria and extracellular space, respectively (34, 35). Therefore, these binding interactions may not be biologically relevant. Additionally, we generated complete titration curves for yeast cells expressing the identified portions of BRD7 and GGNBP2 to estimate their binding affinities for SMAD3 using the multivalent binding model discussed earlier **(Fig. 4B-C)**. The *K*_*D*_ values associated with monovalent bait-prey binding were estimated as 365 nM (309 – 453 nM, 68% confidence interval) and 1165 nM (964 – 1408 nM, 68% confidence interval), respectively for the BRD7 and GGNBP2 fragments **(Fig. 4D)**.

**Figure 4.**
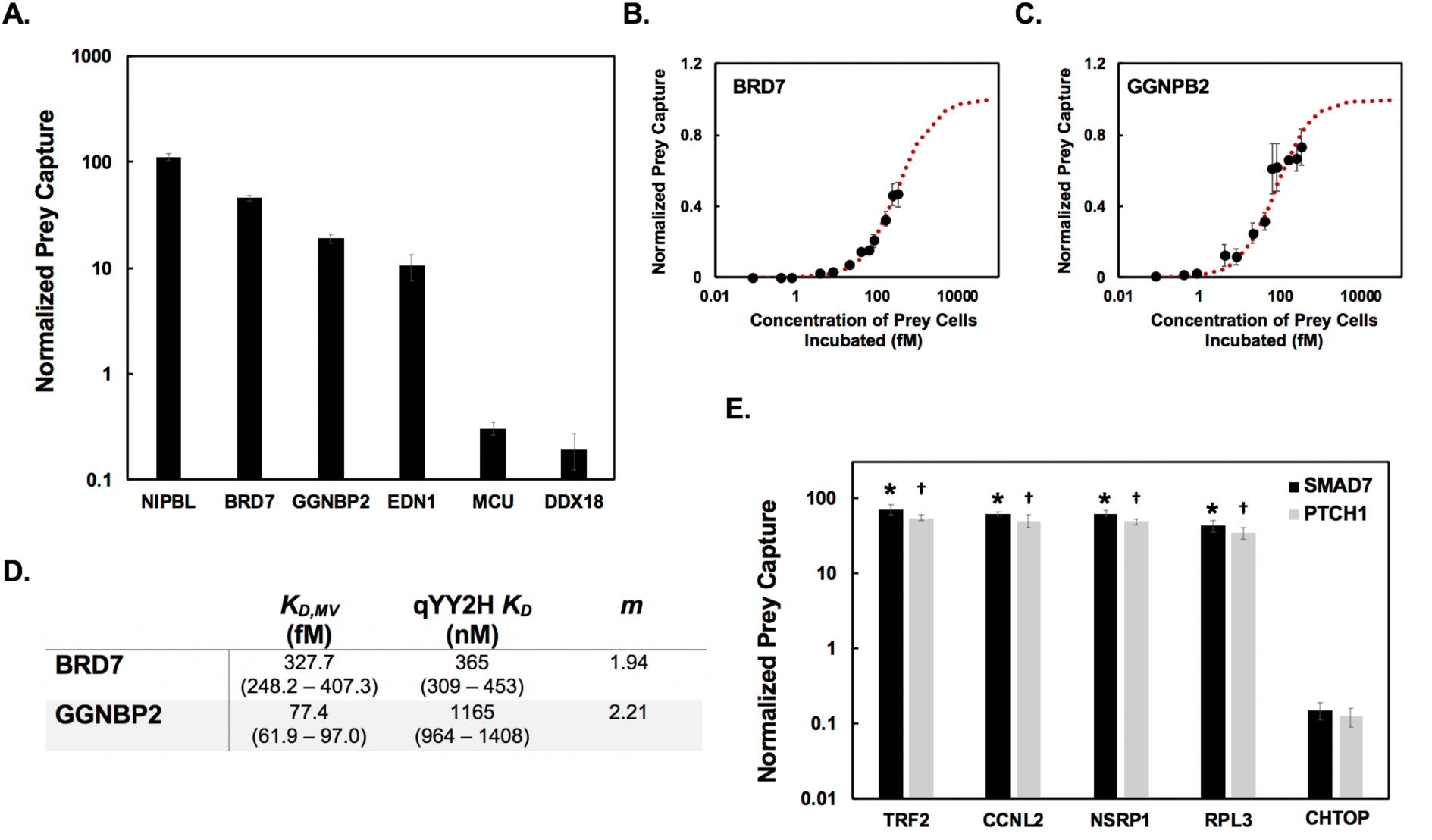
Use of qYY2H to characterize bait-prey pairs identified from a cDNA library screen. For prey isolated from a cDNA library screen, cells co-displaying these prey protein sequences and NanoLuc were incubated with magnetic cells expressing the bait. The luminescence signal of the captured prey cells was used to numerate prey cell capture. **(A)** Prey cell recovery by magnetic bait cells expressing SMAD3 for prey protein sequences identified in cDNA library selections against SMAD3 using qYY2H. Cell capture data for each cDNA prey was normalized by the recovery of cells displaying the SMAD binding domain of SARA. Bait-prey, yeast-yeast titration curves were generated for two of the identified prey, BRD7 **(B)** and GGNBP2 **(C). (D)** A global, non-linear regression was used to estimate *K*_*D,MV*_, the equilibrium dissociation constant describing bait-prey, yeast-yeast binding. Using the previously established semi-empirical model, *K*_*D*_ values (qYY2H *K*_*D*_) describing the binding strength of monovalent bait-prey interactions were estimated as well as associated multiplicity constants, *m*. Values in parentheses represent the bounds of a 68% confidence interval. **(E)** Prey cell recovery by magnetic bait cells expressing the WW domains of YAP using qYY2H. The prey protein sequences considered were identified in a cDNA library screen against the WW domains of YAP. Cell capture data for each cDNA prey was normalized by the recovery of cells displaying either SMAD7 (black) or PTCH1 (grey), known WW binding peptides that contain a PPPY motif. * represents p < 0.05 for a two tailed, paired t-test in comparison to the capture of SMAD7 cells while † represents p < 0.05 for a two tailed, paired t-test in comparison to the capture of PTCH1 cells. Error bars represent the standard error of the mean for three repeats for all data shown.

To further investigate the use of qYY2H in the context of identifying putative PPIs, we screened the cDNA library to identify interaction partners for the tandem WW domains of the YAP protein. The WW domain is a frequently occurring protein-interaction domain that mediates PPIs in signaling pathways by recognizing proline-rich motifs (36–39). Besides proline-rich motifs, WW domains have also been shown to interact with phosphoserine-proline or phosphothreonine-proline motifs (40), LPxY motifs (41), and polyprolines flanked by arginine residues or interrupted by leucine residues (42–44). Using similar methods as described for SMAD3, we identified sequences corresponding to 10 unique proteins that putatively interact with the WW domains of YAP after analyzing 30 clones by DNA sequencing **(Table S2)**. While we did not identify sequences containing the most common PPXY motif, many of the prey identified contained other WW binding motifs. For example, four of the isolated sequences contained a previously identified arginine motif (42). Five sequences contained no previously identified WW binding motifs, consistent with previous cDNA screens to identify WW-binding prey, where at least 20% of the isolated prey did not contain a known binding motif (45).

We further analyzed five of the isolated WW-domain binding sequences using qYY2H **(Fig. 4E)** and compared the strength of their binding interaction with the bait to that of known WW domain binding peptides, SMAD7 and PTCH1. SMAD7 and PTCH1 both contain a PPPY motif, and each binds the WW domains of YAP with *K*_*D*_ = 8 μM and 24.7 μM, respectively (46, 47). Isolated prey containing fragments of the TATA-box binding protein associated factor 2 (TRF2), cyclin L2 (CCNL2), and nuclear speckle splicing regulatory protein 1 (NSRP1) contain an arginine-based binding motif. In contrast, some of the characterized prey, such as sequences from ribosomal protein L3 (RPL3) and chromatin target of PRMT1 protein (CHTOP), do not contain any known WW binding motifs. For the five prey considered, the number of prey cells captured by the WW domain bait cells was greater than the capture of the control cells expressing either SMAD7 or PTCH1, except for the CHTOP prey **(Fig. 4E)**; this suggests that the identified CHTOP sequence likely binds the WW domains of YAP with weaker affinity than the SMAD7 and PTCH1 controls, whereas all other sequences likely bind with similar or higher affinity than SMAD7 and PTCH1.

Collectively, these results show that qYY2H can be used for first discovering putative PPIs by screening cDNA libraries followed by quantitative assessment of the relative binding strengths associated with the identified putative interactions. Importantly, we unveiled a putative binding interaction between the WW domains of YAP and a fragment of CHTOP, with an affinity likely weaker than the YAP-PTCH1 interaction (24.7 μM). Despite the suggested weak affinity of CHTOP for the WW domains of YAP (<24.7 μM), the number of CHTOP-displaying cells captured using the qYY2H assay is well above the detection limit of the assay, as assessed by the lowest number of luciferase-displaying cells quantified using a standard curve in this study. Thus, even weak affinity, putative binding interactions can be identified and quantitatively assessed with qYY2H.

### qYY2H enables evaluation of binding interactions mediated by post-translational modifications

Many processes that regulate biological function rely on protein domains that specifically bind post-translationally modified targets to relay signals (48). For instance, the SH2 and PTB domains bind targets modified by phosphorylation (49). Previous studies have demonstrated the identification of binding interactions dependent on post-translational modifications by linking the modifying enzyme to the bait protein when using the Y2H system (13, 14) or through co-expression of the modifying enzyme in a bacterial two hybrid system (50). We investigated if qYY2H can be extended to quantitatively assess binding interactions mediated by post-translational modifications by utilizing enzymatically modified bait proteins displayed on the yeast surface as previously described (51). In this approach, the modifying enzyme and an Aga2-substrate fusion are co-expressed with ER retention tags to increase their ER residence times. The enzyme acts on the substrate within the ER prior to surface display of the modified Aga2-substrate fusion. Specifically, we evaluated the binding interactions mediated by a phosphopeptide (F1163 – F1183, containing pY1173) derived from epidermal growth factor receptor (EGFR). We used the Abelson tyrosine kinase, which can phosphorylate Y1173 in EGFR, as our modifying enzyme (51, 52). To afford expression of an enzymatically modified bait on the surface of magnetic yeast cells, we refashioned the yeast surface display plasmid to encode the modifying enzyme downstream of the Gal10 promoter **(Fig. S7)**. The Aga2 fusions containing the substrate peptide and the iron-oxide binding protein SsoFe2 are encoded downstream of a Gal1 promoter on the same plasmid. The Aga2-substrate peptide and the Aga2-SsoFe2 fusions are translated as separate proteins through the inclusion of a T2A ribosomal skipping peptide as we have previously described (19). Flow cytometry analysis using an anti-phosphotyrosine antibody showed that the EGFR peptide containing Y1173 is phosphorylated only in the presence of Abelson kinase **(Fig. 5A)**.

**Figure 5.**
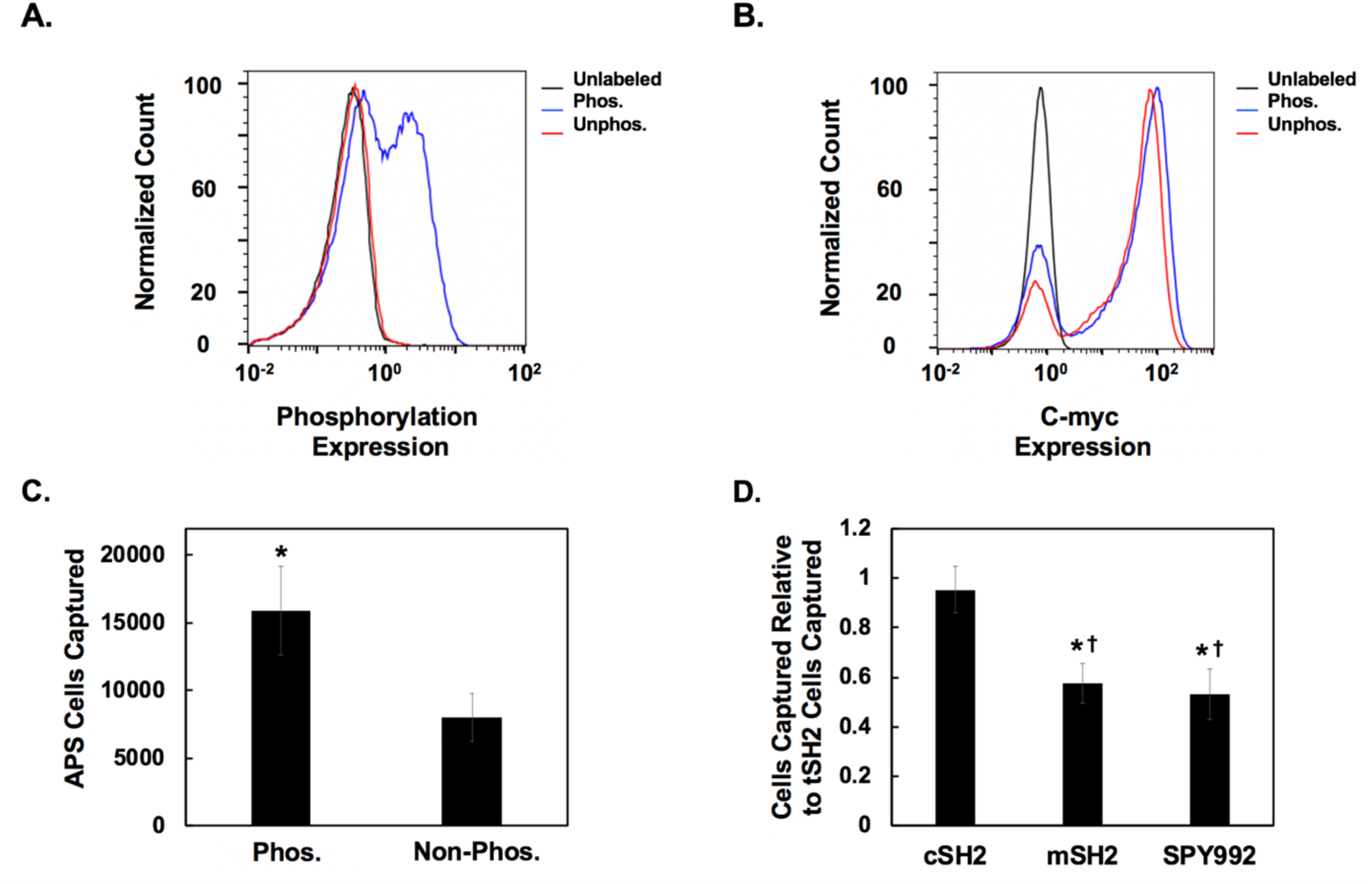
Use of qYY2H system to evaluate protein-protein interactions that depend on a post-translationally modified binding domain. **(A)** Yeast-cells expressing a phosphorylated EGFR peptide (pY1173) were used as bait. The EGFR peptide was enzymatically modified within the ER by Abelson tyrosine kinase prior to surface display. Flow cytometry analysis was used to evaluate the fraction of the bait population expressing the phosphorylated peptide above background. Yeast cells containing the modifying kinase (blue) as well as non-enzymatically modified yeast cells (red) were labeled with an anti-phosphotyrosine Alexa Fluor 647 antibody followed by flow cytometry detection. The fluorescence of unlabeled cells (black) is also considered. **(B)** Peptides may be displayed prior to modification by the kinase. Hence, a fraction of cells from the population that co-expresses Abelson kinase and the substrate peptide will display non-phosphorylated peptides. To quantify total peptide display, regardless of phosphorylation state, the cell population containing the modifying kinase (blue) as well as the non-enzymatically modified yeast (red) were labeled with an anti-c-myc antibody followed by detection using a goat-anti-chicken 488 antibody and flow cytometry analysis. The fluorescence of unlabeled cells (black) is also considered. Representative flow cytometry plots from three independent repeats are shown. **(C)** Yeast cells co-displaying the SH2 domains of adaptor protein APS and NanoLuc were incubated with magnetic bait cells expressing either an EGFR peptide phosphorylated at Y1173 or an unphosphorylated EGFR peptide. The cell capture values associated with the phosphorylated bait were normalized by the fraction of bait cells expressing c-myc fusions that were also phosphorylated. A greater number of APS prey cells were recovered by the phosphorylated bait cells than the non-phosphorylated bait cells. * represents p < 0.05 for a two-tailed, paired t-test in comparison to the non-phosphorylated bait. Error bars represent the standard error of the mean for seven repeats. **(D)** Yeast cells co-expressing tSH2, cSH2, mSH2, or SPY992 and NanoLuc were incubated with magnetic bait yeast expressing the pY1173 EGFR peptide. Cell capture values for each prey were first normalized by the fraction of bait cells expressing c-myc fusions that were also phosphorylated. After, the normalized cell capture values for each repeat were compared to the normalized capture of tSH2 cells by the phosphorylated bait cells. * represents p< 0.05 for a two-tailed, paired t-test in comparison to the capture of tSH2 cells while † represents p< 0.05 for a two-tailed, paired t-test in comparison to the capture of cSH2 cells. Error bars represent the standard error of the mean for at least five repeats.

To assess the use of qYY2H for studying phosphorylation-dependent binding, we first evaluated the interaction between the SH2 domains of the adapter protein APS and pY1173. Previous cDNA library screens identified that the SH2 domains of adapter protein APS bind to EGFR pY1173 (53). APS was shown to not interact with the non-phosphorylated EGFR peptide. Prey cells expressing the SH2 domains of APS and NanoLuc were incubated with magnetized bait cells expressing the phosphorylated EGFR peptide pY1173 obtained by enzymatic modification by Abelson kinase within the ER, or the non-phosphorylated peptide as a control. Subsequently, bait-prey complexes were separated using a magnet and the number of prey cells captured was quantified. A fraction of cells from the population that co-expresses Abelson kinase and the substrate peptide will display non-phosphorylated peptides. Hence, the cell capture data for the bait population co-expressing Abelson kinase was normalized by the fraction of cells expressing peptide fusions that are also phosphorylated as evaluated by immunofluorescent detection **(Fig. 5A-B)**. A greater number of APS displaying prey cells was captured when the phosphorylated peptide was used as bait **(Fig. 5C)**, suggesting the qYY2H can be used to identify interactions that rely on post-translational modifications.

We further investigated if qYY2H could be used to quantitatively discriminate between proteins that bind a post-translationally modified target with varying interaction strengths. SH2 domains bind phosphotyrosine residues promiscuously via a consensus sequence pYXXP motif (54–57). The tandem SH2 domains of phospholipase C–γ1 (PLCγ1), denoted tSH2, bind to phosphorylated EGFR. tSH2, as well as its C-terminal SH2 domain, denoted cSH2, have been evaluated as biosensors for live cell imaging of EGFR phosphorylation at Y992 (58–60). However, both tSH2 and cSH2 show promiscuous binding to other phosphorylated residues in EGFR (58, 61). To overcome the promiscuity of tSH2 and cSH2, Tiruthani *et al*. identified two proteins with enhanced specificity for pY992, mSH2 and SPY992, which were isolated by screening combinatorial libraries generated by mutagenesis of cSH2 and a Sso7d protein scaffold, respectively (58). SPY992 and mSH2 exhibit high specificity of binding to pY992 over other phosphorylation sites in EGFR, relative to tSH2 and cSH2. We used qYY2H to investigate binding of tSH2, cSH2, mSH2, and SPY992 to pY1173. Consistent with lower binding of mSH2 and SPY992 to sites other than pY992, fewer prey cells expressing mSH2 and SPY992 were captured by bait expressing pY1173 than cells expressing tSH2 and cSH2 as prey **(Fig. 5D)**. These results show that qYY2H can be used to quantitatively compare binding interactions that are dependent on post-translational modification of the bait protein.

In conclusion, we have developed a quantitative yeast-yeast two-hybrid system, denoted qYY2H, wherein the bait and prey proteins are expressed as yeast cell surface fusions. qYY2H can be used not only to screen cDNA libraries to identify PPIs, but also to quantitatively assess the strength of bait-prey binding interactions. Notably, PPIs dependent on post-translational modification of the bait can be analyzed using qYY2H. Due to the multivalent nature of the interaction between yeast displayed bait and prey, qYY2H can be used to investigate binding interactions with low binding affinities. In this study we have identified binding interactions with *K*_*D*_ ∼ at least 25 μM, where the signal was well above the limit of detection of qYY2H; therefore, we expect that interactions with weaker affinities may be assessed. Further, we have shown that the strength of the multivalent interaction between bait and prey yeast bears a quantitative relationship with the binding affinity of the monovalent bait-prey interaction (*K*_*D*_). We have described a mathematical framework to exploit this relationship for quantitative estimates of monovalent bait-prey interaction *K*_*D*_. We anticipate that qYY2H will be a powerful tool for quantitative analysis of protein-protein interactions. In particular, we expect that qYY2H will be very useful for efficient characterization of binding proteins isolated from combinatorial libraries. Finally, the quantitative framework discussed herein for yeast-yeast binding interactions will be useful for designing combinatorial screens for isolating binders from yeast display libraries using whole cell targets (62) or when the target protein is expressed as a yeast cell surface fusion (19).

## Materials and Methods

### Plasmid construction for dual display of prey and NanoLuc

The previously described pCT302-SsoFe2-T2A-TOM22 plasmid was altered to afford dual expression of a prey protein and NanoLuc using a ribosomal skipping T2A peptide (19). DNA encoding a prey protein was inserted between the NheI and BamHI sites while DNA encoding NanoLuc was inserted between the AvrII and NdeI sites. Detailed cloning protocols are provided in the supplemental methods.

### Luciferase-based binding quantification assays using bait expressing magnetic yeast

Yeast cells co-expressing the bait protein and SsoFe2 were magnetized and blocked as previously described (19). Briefly, 1×10^7^ magnetic bait cells were incubated with varying concentrations of yeast cells co-expressing the prey and NanoLuc in addition to 1×10^9^ EBY100 cells in 2 mL of 0.1% PBSAT (PBS pH 7.4, 0.1% BSA, 0.05% tween-20) for 1 hour. When comparing the binding affinity of mutants for a signal prey concentration, only 1×10^7^ prey cells were incubated. When generating titration curves to estimate binding affinity values, the number of prey cells incubated ranged from 1×10^5^ - 4×10^8^ cells. After, any cells not bound to the magnetic bait cells were removed using a magnet. The magnetic bait cells and any bound prey cells were washed 3X with 0.1% PBSAT and then resuspended in 100 μL of PBS. The Nano-Glo Luciferase Assay system (Promega) was used to detect binding of prey cells to the immobilized bait proteins. 100 μL of the Nano-Glo reagent was added to the magnetic bead solution. The reaction proceeded for 3 minutes before placing the tube onto the magnet and plating 100 μL of the reaction in duplicate onto a 96 white-well plate with a clear bottom. The luminescence was read using a Tecan Infinite 200 plate reader using an integration time of 1000 ms, settle time of 0 ms, and no attenuation. Additional details are provided in the supplemental methods.

A calibration curve was made for each prey construct to develop a relationship between luminescence signal and the number of prey cells present. Luminescence assays were carried out as previously described to determine the points in the calibration curve. The curve was generated using 1×10^3^ - 1×10^6^ prey cells resuspended in 100 μL of PBS. The luminescence of just PBS and the Nano-Glo reagent was also measured as a blank. The calibration curves were generated by plotting background subtracted luminescence signals vs number of cells and fitting a linear regression. Accordingly, the generated calibration curve was used to predict the exact number of prey cells captured by the magnetic bait cells by relating the luminescence signal produced by the captured prey cells.

### Luciferase-based binding quantification assays using bait functionalized magnetic beads

25 μL of Dynabeads Biotin Binder Beads (Thermo Fisher Scientific) were functionalized overnight with biotinylated, soluble bait protein (10.8 μM biotinylated lysozyme or 0.1 μM biotinylated IgG). The following morning the beads were washed 3X with 0.1% PBSA (PBS pH 7.4, 0.1% BSA) followed by blocking in 1 mL of 1% PBSA (PBS pH 7.4, 1% BSA) for two hours. After, the beads were incubated with varying concentrations of yeast cells co-expressing the prey protein and NanoLuc along with 1×10^9^ EBY100 cells in 2 mL of 0.1% PBSAT for 1 hour at room temperature. After, any cells not bound to the bait-functionalized magnetic beads were removed using a magnet. The beads were washed 3X with 0.1% PBSAT and then resuspended in 100 μL of PBS. The Nano-Glo Luciferase Assay was performed as previously described to detect binding of the prey cells to the bait-functionalized magnetic beads. Subsequently, a luminescence calibration curve was used to quantify the number of prey cells captured. Additional details are provided in the supplemental methods.

### Applying Non-linear Regression Model to Estimate K_D,MV_

Titration curves detailing the binding of prey cells to magnetic bait cells or magnetic beads were generated using the previously described binding assays. The number of prey cells (1×10^5^ - 4×10^8^ cells) incubated was varied while the number of magnetic bait cells or magnetic beads was held constant (1×10^7^ cells or beads). A monovalent binding isotherm, as described in Equation 1, was applied to the data to predict *K*_*D,MV*_ using a global fit. When fitting the isotherm, some data points were not used due to non-specific binding or the Hook effect. Details on how points were selected for fitting **Equation 1** are described in the supplemental methods.

### Construction of a yeast-displayed cDNA library with a luciferase reporter

A yeast displayed cDNA library that concurrently displays NanoLuc (diversity ∼6×10^6^) was constructed using DNA amplified from the Clontech Mate & Plate Library-Universal Mouse (Normalized) cDNA library (Takara Bio Inc). The yeast library was generated using the previously described lithium acetate yeast transformation method (63). A detailed protocol is described in the supplemental methods.

### Screening a yeast-displayed cDNA library with a luciferase reporter to discover putative PPIs

Two rounds of magnetic sorting were carried out to isolate putative binders to SMAD3 and the WW domains of YAP. In these sorts, the bait protein was displayed on the surface of magnetic yeast cells. For the first screening round, 5×10^7^ magnetic bait cells were prepared as previously described followed by incubation with 1×10^8^ induced cDNA, NanoLuc library cells and 1×10^9^ EBY100 cells in 2 mL of 0.1% PBSAT for 2 hours at room temperature. Any library cells bound to the magnetic bait cells were isolated using a magnet followed by washing with 0.1% PBSAT (5X) prior to expansion in 20 mL of SDCAA (-TRP media).

In the second magnetic screening round, 5×10^6^ magnetic bait cells were incubated with 1×10^7^ library cells from the first screening round and 1×10^9^ EBY100 cells. The library cells bound to the magnetic bait cells were expanded in 5 mL of SDCAA (-TRP) media after washing. DNA was recovered from 30 individual clones using a Zymoprep yeast plasmid miniprep II kit (Zymo Research) and sequenced. Additional details can be found in the supplemental methods.

## Supporting information

Glucose oxidase and horseradish peroxidase reporter assay results, supplementary materials and methods, strategy for quantitative yeast-yeast two hybrid assay, evaluation of GOx as a reporter, bait-prey yeast-yeast titration curves for NB.TOM22 and Sso7d.TOM22.2, comparison of bait protein expression levels, relationship between monovalent *K*_*D*_ and multivalent *K*_*D,MV*_ for yeast-magnetic bead system, comparison of *K*_*D,MV*_ for yeast-yeast interactions and *K*_*D,MV*_ for yeast-bead interactions, plasmid schematic for co-expression of enzyme modified bait and iron oxide binding protein, qYY2H titration curves along with slope plots, a table detailing the putative interacting prey identified for SMAD3, a table detailing the putative interacting prey identified for the WW domains of YAP, a table of gene block fragments, and a table of oligonucleotide primers.

## Acknowledgments

This work was funded by two grants from the National Science Foundation (CBET 1511227 and CBET 1510845). KB kindly acknowledges support from an NSF Graduate Research Fellowship. KB and JB kindly acknowledge support from a National Institute of Health Molecular Biotechnology Training Fellowship (NIH T32 GM008776).

