## Supporting Information for "Quantitative yeast-yeast two hybrid for discovery and binding affinity estimation of protein-protein interactions"

**This PDF file includes:**

Supplementary Results and Discussion
Supplementary Materials and Methods
Figures S1 to S19
Tables S1 to S4

**Supplementary Results and Discussion**

*Enzyme sensitivity is important to quantitatively assess bait-prey interactions.*

Multiple reporter enzymes were evaluated for their sensitivity to distinguish the binding interactions between different prey for a given bait. The first enzyme considered was glucose oxidase (GOx), which uses a colorimetric assay for detection. First, we incubated yeast cells co-expressing prey proteins and GOx with magnetic beads functionalized with the soluble bait protein. Any prey cells bound to the magnetic beads were isolated and incubated with the GOx assay reagent. The assay reagent contains dextrose, 3,3′,5,5′-Tetramethylbenzidine (TMB), and horseradish peroxidase (HRP). In the presence of GOx, glucose reacts with oxygen to form glucono delta-lactone and hydrogen peroxide. HRP reduces the generated hydrogen peroxide to water using TMB as a hydrogen donor, resulting in the formation of a diimine that causes the solution to turn blue. The reaction is halted by the addition of sulfuric acid, resulting in the solution turning yellow. The intensity of the color change can be read using a spectrophotometer. The intensity of the color change is directly proportional to the concentration of GOx present in the solution and, accordingly, the number of prey cells captured.

Three previously identified prey, derived from the Sso7d scaffold, with affinity for lysozyme were considered, Sso7d.BVL.Lys ($K_D$=1.7 nM), Sso7d.CTL.Lys ($K_D$=250 nM), and Sso7d.NTL.Lys ($K_D$=1300 nM) as well as an irrelevant Sso7d mutant, Sso7d.hFc (1, 2). The GOx signal was higher for each of the lysozyme mutants in comparison to the negative control, suggesting that the bait functionalized magnetic beads were capturing cells expressing the lysozyme binding proteins at a higher level than cells expressing the irrelevant Sso7d mutant. However, the difference in signal between the lysozyme binding prey was not significantly different, though the mutants vary in affinity. These results suggest that GOx as a reporter is not sensitive enough to distinguish the affinities between different mutants. GOx can be used to distinguish if two proteins interact, but it cannot be used to rank order affinities between different prey proteins for a single bait protein. The sensitivity of the assay may increase if the TMB substrate concentration is higher. However, the solubility of the TMB substrate was reached in these assays (3).

We also evaluated the use of GOx as a reporter when the bait protein is expressed on the surface of magnetic yeast cells (4). For these assays, three previously identified prey, derived from the Sso7d scaffold, with affinity for the Fc portion of human IgG (hFc), Sso7d.hFc ($K_D$=450 nM),

Sso7d.his.hFc ($K_D$=1647 nM), and Sso7d.ev.hFc ($K_D$=5280 nM), were co-expressed with GOx (2). An irrelevant Sso7d mutant, Sso7d.CTL.Lys, was also considered (1). Here, the GOx assays performed on the captured prey cells barely generated a signal above background. It is important to note that the prey cells were not removed from the magnetic bait cells prior to performing the GOx assay. Color formation depends on the interaction of hydrogen peroxide with HRP which could be negatively affected if the generated hydrogen peroxide decomposes in the presence of the iron oxide used for magnetization instead of interacting with HRP (5–7). Iron oxide has also been shown to promiscuously bind to many proteins in solution (8–12). If HRP were to associate with iron oxide, thereby blocking the substrate binding site, then the assay's productivity would be negatively affected.

Subsequently, HRP was explored in a similar manner for its use as a reporter enzyme due to the limitations observed with GOx. HRP activity was detected using 1-Step Ultra TMB Elisa Substrate solution. However, it is likely the substrate solution is lysing any yeast cells present due to its low pH (3.35-3.75). After lysis, any peroxidases from inside the cell are released, compounding the signal readout beyond the signal associated with surface displayed HRP. We tested this hypothesis by incubating non-displaying EBY100 cells with the substrate solution. We were able to obtain a signal even though these cells do not express surface displayed HRP, suggesting that the cells are lysed after the substrate solution is added.

### Supplementary Materials and Methods

*Yeast culturing and plasmids*

*Saccharomyces cerevisiae* strain EBY100 cells containing the pCTCON plasmid were cultured using SDCAA (-Trp) media while cell surface protein expression was induced using SGCAA (-Trp) medium, as previously described (13). Cells containing the pCT302 plasmid were cultured and induced with SDSCAA (-Leu) and SGSCAA (-Leu) media, which is similar in composition to SDCAA and SGCAA media except it contains a synthetic drop out mix (1.62 g/L; US Biological Life Sciences) lacking leucine instead of casamino acids. Cells were grown overnight in SDCAA or SDSCAA media at 30°C with shaking at 250 RPM. Protein expression was induced at 20°C with shaking at 250 rpm after transferring the yeast cells into SGCAA or SGSCAA medium at an OD of 1. EBY100 cells without plasmid were grown in YPD medium (10.0 g/L yeast extract, 20.0 g/L peptone, and 20.0 g/L dextrose) at 30°C with shaking at 250 rpm. Plasmid DNA was transformed using chemically competent EBY100 generated using the Frozen-EZ Yeast Transformation Kit II (Zymo Research).

*Plasmid construction for dual display of prey and reporter enzyme*

Plasmids were constructed to afford the co-expression of a prey protein and a reporter enzyme as yeast cell surface fusions. Previously described plasmids that confer dual protein expression via a ribosomal skipping T2A peptide were modified (4). Specifically, the pCT302-SsoFe2-T2A-TOM22 plasmid was altered by digesting between the AvrII and NdeI sites. DNA encoding specific reporter enzymes, including glucose oxidase (GOx), horseradish peroxidase (HRP), and NanoLuc, were inserted between these sites to generate different dual enzyme display plasmids, pCT302-T2A-GOx, pCT302-T2A-HRP, and pCT302-T2A-NanoLuc, respectively. The GOx gene was amplified using primers Pf1 and Pr1 from genomic DNA extracted from *A.niger* spores as described in Cruz *et al* (14). DNA encoding horseradish peroxidase (Q1 – S308, assuming no signal peptide) was amplified from gene block 1 using primers Pf2 and Pr2 (15). Similarly, DNA encoding the NanoLuc protein was amplified using gene block 2 and primers Pf3 and Pr3 (16). Previously identified binding proteins evolved from the Sso7d scaffold were inserted into each of these plasmids via the NheI and BamHI sites. Sso7d.hFc, Sso7d.his.hFc, and Sso7d.ev.hFc were amplified using primers Pf4 and Pr4 from plasmids described in Gera *et al* (2). Similarly, DNA encoding SSo7d.NTL.Lys and Sso7d.BVL.Lys was amplified from plasmids described by Carlin *et al*. with primers Pf4 and Pr4 (1) while Sso7d.Lys.CTL was amplified with primers Pf5 and Pr4 (1).

A plasmid, pcTCON-NanoLuc, was also constructed that solely expresses NanoLuc as an Aga2-yeast surface fusion. This plasmid was created by amplifying gene block 2 with primers Pf6 and Pr6 and inserting this product between the NheI and BamHI cut sites in pCTCON.

Double stranded gene fragments were purchased from Integrated DNA technologies (IDT). Primer oligonucleotides were purchased from IDT or Eton Biosciences. Gene fragment and primer sequences can be found in **Tables S3-S4**. PCR reactions using Phusion Polymerase (Thermo Fisher Scientific) were performed in a 50 μL volume following the manufacturer's protocols. Restriction enzymes were purchased from New England Biolabs. Restriction digestions of plasmids and PCR products were performed for 4 hours at 37°C in 50 μL reactions with a 5X excess of each restriction enzyme. Digested plasmids were treated for 1 hour at 37°C with Antarctic phosphatase (New England Biolabs). Subsequently, digested products were purified using the 9K series gel and PCR extraction kit (BioBasic). Ligations of digested plasmids and PCR product inserts were performed overnight using T4 DNA ligase (Promega) at 16°C. Ligations were transformed into electrocompetent Novablue *E.coli* cells. Each electroporation was performed at 1.6 KV, 25 μF, and 200 Ω. Plasmid was harvested from overnight *E.coli* cultures using the GeneJET plasmid miniprep kit (Thermo Fisher Scientific).

*GOx-based binding quantification assays using bait-functionalized magnetic beads*

Magnetic beads were functionalized with lysozyme by incubating 25 μL of Dynabeads Biotin Binder Beads (Thermo-fisher Scientific) with 10.8 μM biotinylated lysozyme overnight at 4°C. The beads were washed 2 times with 1% PBSA (PBS pH 7.4, 1% BSA) the next day and blocked with 1 mL of 1% PBSA for 3 hours at 4°C. Next, the magnetic beads were washed two times with 0.1% PBSA (PBS pH 7.4, 0.1% BSA) followed by incubation with $1x10^8$ cells co-expressing the prey protein and GOx along with $1x10^9$ EBY100 cells in 2 mL of 0.1% PBSA for 2 hours. The prey proteins considered were Sso7d.BVL.Lys, Sso7d.CTL.Lys, and Sso7d.NTL.Lys as well as an irrelevant protein, Sso7d.hFc. After, the incubations were placed onto a magnet to isolate any cells bound to the magnetic beads. The magnetic beads and any associated prey cells were then washed 3X with PBS. The beads were resuspended in 100 μL of PBS prior to adding 100 μL of the 2X TMB solution (0.1 mg/mL TMB, 0.4 M dextrose, 1.5 μg/mL HRP diluted in

PBS). This reaction took place in the dark for 1 hour at room temperature before it was quenched with 50 μL of 2 M sulfuric acid. The reaction tubes were placed onto a magnet, and 100 μL aliquots were transferred onto a 96 well plate in duplicate. Each well's absorbance was measured at 450 nm using a plate reader.

The cell surface expression level of the co-displayed prey and GOx proteins was estimated using immunofluorescence detection. The expression of the prey protein was detected using a chicken anti-c-myc antibody (Thermo Fisher Scientific) while the expression of the GOx protein was detected using a mouse anti-V5 antibody (Thermo Fisher Scientific). To begin, $5x10^6$ cells were labeled with a 1:100 dilution of the primary antibodies for 15 minutes at room temperature. After, a secondary labeling was performed using a 1:250 dilution of goat-anti-chicken 488 or a donkey anti-mouse 633 antibody (Immunoreagents) for 10 minutes on ice. All labelings were conducted in 50 μL of 0.1% PBSA. Washes took place between each labeling. The binding of the antibodies was detected using a Miltenyi Biotec MACsQuant VYB cytometer.

An activity control was also carried out for each prey construct. Briefly, $1x10^6$ cells co-expressing the prey and GOx were washed two times with PBS and ultimately resuspend in 100 μL of PBS prior to the addition of 100 μL of the 2X TMB solution. The control reactions were allowed to proceed for 5 minutes prior to being quenched with 50 μL of sulfuric acid. The quenched reactions were centrifuged. 100 μL aliquots of the supernatant plated in duplicate were read at 450 nm using a plate reader as previously described.

The background absorbance associated with an equal volume mixture of PBS and the TMB solution at 450 nm was subtracted from each raw, experimental absorbance value. For each prey, the background subtracted absorbance value associated with bait-prey binding was normalized by first dividing by the prey cells' mean fluorescence of c-myc expression and then dividing by the background subtracted absorbance of that particular prey's activity control. These normalizations were performed to account for differences in prey and GOx expression levels, respectively.

The lysozyme protein used as the bait was biotinylated by incubating a 3:1 molar excess of Ez-Link™Sulfo-NHS-LC-Biotin (Thermo-Fisher Scientific) for two hours at 4°C. Excess reagent was removed through dialysis against 50 mM Tris-HCl, 300 mM NaCl pH 7.5. A BCA assay was used to estimate the protein concentration.

*GOx-based binding quantification assays using bait expressing magnetic yeast*

Yeast cells co-expressing hFc and SsoFe2 were previously generated and used as bait (4). The bait cells were magnetized and blocked as previously described (4). Briefly, $1 \times 10^8$ cells co-expressing the bait protein and SsoFe2 were incubated with 2 mL of an iron oxide solution (4 mg/mL iron oxide in water) in a total of 10 mL of 1% TBSA (50 mM Tris, 300 mM NaCl pH 7.4, 1% BSA) for 20 minutes at room temperature. Non-magnetized cells were removed using a magnet. $OD_{600}$ of the cells prior to the addition of the iron oxide ($OD_i$) and after the removal of yeast bound to iron oxide ($OD_f$) was measured using 100 μL of the sample. The number of cells bound to the iron oxide was calculated as ($OD_i$-$OD_f$). The magnetized cells were washed 3X with 1% TBSA and then aliquoted for use in each assay ($1 \times 10^7$ magnetic bait cells per assay). The aliquoted magnetized bait cells were then blocked for 1 hour using 2 mL of 1% TBSA and then washed two more times with 0.1% TBSA (50 mM Tris, 300 mM NaCl pH 7.4, 0.1% BSA). $1 \times 10^7$ magnetic target cells were incubated with $5 \times 10^8$ yeast cells co-expressing the prey protein and GOx along with $1 \times 10^9$ EBY100 for 2 hours at room temperature in 2 mL of 0.1% TBSA. The incubations were then placed onto a magnet, and unbound cells were removed. The magnetic bait cells and any bound prey cells were washed 3X with 0.1% PBSA prior to being resuspended in 100 μL of PBS. 100 μL of the 2X TMB solution was subsequently added. The reaction proceeded for 1 hour in the dark prior to quenching with 50 μL of sulfuric acid. The reactions were placed onto a magnet and a similar process, as previously described for the magnetic beads, was used to measure the absorbance of the reactions. Activity control assays and flow cytometry analysis of the prey cells were completed as previously described for normalization.

*Magnetization of bait expressing cells for NanoLuc reporter assays*

Yeast cells co-expressing a bait protein (hFc or TOM22) and SsoFe2 were magnetized in a similar manner as for the yeast-yeast GOx assays. These bait cells were previously generated in Bacon *et al* (4). Here, PBS based buffers were used instead of Tris based buffers. Specifically, the bait cells were magnetized in 1% PBSA. Similarly, $1 \times 10^7$ magnetized target cells were used in each assay. After aliquoting, the magnetized target cells were blocked with 2 mL of 1% PBSA for 1 hour at room temperature. Following blocking, the magnetic target cells were washed 2X and resuspended with 0.1% PBST (PBS pH 7.4, 0.1% BSA, 0.05% tween-20) prior to incubation with the prey cells.

*Semi-empirical framework to estimate $K_D$ using qYY2H*

Using the qYY2H system, titration curves were generated from bait-prey binding assays using luminescence quantification as described. $1x10^7$ magnetic bait cells were used for each assay while the number of the number of prey cells varied from $1x10^5$ cells - $4x10^8$ cells. Also included were $1x10^9$ EBY100 cells. The Nano-Glo assay described in the main text was used to detect binding of the prey cells to the magnetic bait cells. A luminescence standard curve was used to predict the number of prey cells captured as described in the main text. Three or four repeats were completed for each bait-prey pair. Prey considered with affinity for hFc included Sso7d.hFc, Sso7d.his.hFc, and Sso7d.ev.hFc (2). Prey considered with affinity for TOM22 included Sso7d.TOM22.1, Sso7d.TOM22.2, NB.TOM22.1, and NB.TOM2.2 (4).

The number of prey cells incubated and subsequently captured were converted to a concentration using the following relationship:

$$[Cell] = \frac{N}{N_{AV}V} \quad \textbf{(Equation S1)}$$

Where $[Cell]$ is the concentration of cells in moles, $N$ is the number of cells, $N_{AV}$ is Avogadro's number, and $V$ is the volume the assay took place in (i.e.: 2 mL).

Curves were generated comparing the concentration of prey cells captured vs the concentration of prey cells initially incubated. The number of bait-prey, yeast-yeast complexes isolated by magnetic separation was related to the apparent affinity of multivalent interaction between bait and prey yeast ($K_{D,MV}$) using a monovalent binding isotherm:

$$[Complex] = \frac{[Max][Prey]}{K_{D,MV}+[Prey]} \quad \textbf{(Equation S2)}$$

Where $[Complex]$ is the molar concentration of bait-prey, yeast-yeast complexes, $[Prey]$ is the initial molar concentration of prey yeast used in the experiment, and *[Max]* is the maximum molar concentration of prey cells that are captured for a given number of magnetized bait yeast. Typically, all data points in the titration curve were used when fitting the monovalent isotherm. However, some data points in the titration curves for individual repeats were not included when fitting the non-linear regression to estimate *[Max]* and $K_{D,MV}$ values due to non-specific binding or observation of the Hook effect. When evaluating non-specific binding, the slope between adjacent data points was calculated and plotted against the concentration of prey cells incubated. When generating this plot, the larger of the incubated prey cell concentrations, among the data points, was used as the x coordinate. For a given replicate, the first four slope points were ignored. When the slope reached ~0 (generally <0.005), we assumed that binding was reaching saturation. Sometimes multiple instances of a slope ~0 would occur for a particular plot. In these instances, we chose to not include data that occurred for concentrations higher than the second instance of slope ~0 to eliminate bias if the first minimum slope occurred due to error. For some prey, the slope increased, after reaching a minimum, therefore this data was not included when fitting the monovalent binding isotherm, as we assumed this increase in binding was due to non-specific binding. Data was also not included if the number of cells captured was significantly greater than the number of cells captured at the neighboring concentrations of incubated prey cells. For yeast-yeast binding, **Figs. S8-S14** describe the data used when fitting the monovalent binding isotherm to the generated titration curves for each repeat as well as the slope plots used to determine what data to include. For each bait-prey interaction, a global fit was used to estimate [*Max*] values for each repeat and a single $K_{D,MV}$ across the repeats by minimizing the sum of the squared error.

It was assumed that the apparent affinity of the multivalent interaction between the bait and prey yeast ($K_{D,MV}$) can be related to the binding affinity of the monovalent bait-prey interaction ($K_D$) by

$$K_{D,MV} = {K_D}^{m} \quad \textbf{(Equation S3)}$$

where *m* is a multiplicity constant that captures the increase in apparent affinity due to multivalent interactions. For each bait-prey considered, estimates for $K_D$ had been previously predicted by performing yeast surface titrations using soluble protein. We used the fitted values of $K_{D,MV}$ and the previously estimated $K_D$ values to calculate *m* for five of the bait-prey pairs. We used a random number generator to choose which interaction prey to consider (TOM22.NB.1, Sso7d.hFc, Sso7d.TOM22.1, Sso7d.his.hFc, Sso7d.ev.hFc). After, we generated a plot of comparing *m* and $K_D$ for the interaction pairs considered. This data fit a power law relationship where

$$m = 10.277{K_D}^{0.1125} \quad \textbf{(Equation S4)}$$

This relationship can be inserted into **Equation S3** to predict $K_{D,MV}$ as a function of $K_D$ as follows

$$K_{D,MV} = (K_D)^{10.277*{K_D}^{0.1125}} \quad \textbf{(Equation S5)}$$

We used estimates of $K_{D,MV}$ for the other considered prey, Sso7d.TOM22.2 and NB.TOM22.2, to predict values of $K_D$ using **Equation S5** to evaluate the accuracy of our developed semi-empirical model.

A relationship between [*Max*] and $K_D$ was also established for the five bait-prey pairs randomly chosen above. For each repeat, the [*Max*] value estimated when applying the monovalent binding isotherm to the titration curve data was normalized by dividing by the average concentration of prey cells captured for the data included when applying the non-linear regression model. If a particular piece of data was not included when fitting the monovalent binding isotherm, then it was not used to calculate this cell capture average. For a given interaction pair, normalized [*Max*] values were averaged across the repeats. A plot was generated of the averaged, normalized [*Max*] values vs $K_D$ and a power law regression was applied. $K_D$ values were previously estimated for these bait-prey pairs using yeast surface titrations of soluble protein.

A similar procedure was used to generate titration curves when soluble bait protein was immobilized onto magnetic beads. We only considered human IgG (hFc) as a bait when evaluating this system using the NanoLuc reporter, but it could be extended to other soluble bait proteins. First, 25 μL of Dynabeads Biotin Binder Beads were washed two times with 0.1% PBSA. After, the beads were incubated overnight with 100 μL of 0.1 μM biotinylated human IgG (Immunoreagents) diluted in 0.1% PBSA based on the manufacturer's recommendations. The next day the beads were washed 3X with 0.1% PBSA followed by blocking in 1 mL of 1% PBSA for two hours. After, the beads were washed an additional two times in 0.1% PBSA prior to incubation with the prey cells and $1 \times 10^9$ EBY100 in 2 mL of 0.1% PBSAT. The number of prey cells incubated varied ranging from $1 \times 10^5$ cells - $4 \times 10^8$ cells. After separation of unbound cells and washing (3X with 0.1% PBSAT), the Nano-Glo assay was used to detect binding of the prey cells to the magnetic beads as described in the main text. Standard curves generated using known quantities of prey cells were used to estimate the number of prey cells captured. Three repeats were repeated for each prey. The monovalent binding isotherm as described in **Equation S2** was fit to this data, in a similar manner as previously described, to estimate [*Max*] values for each repeat and a single $K_{D,MV}$ across all repeats for each prey. Previously described methods were used to determine which points to include when applying the fit **(Fig. S15-S17)**. Plots of $m$ vs $K_D$ and normalized [*Max*] vs $K_D$ were generated and fit using a power law regression as previously described **(Fig. S5)**.

*Construction of a yeast displayed cDNA library with a luciferase reporter*

The Clontech Mate & Plate Library- Universal Mouse (Normalized) cDNA library was obtained from Takara Bio Inc. The library was grown in SDSCAA (-Leu) media at 30°C. Plasmids were extracted from these cells using a Zymoprep Yeast Plasmid Miniprep Kit II (Zymo Research).

Twenty four plasmid extractions using $9x10^6$ cells were performed. The prey DNA was amplified using primers Pf7 and Pr7 in 24 identical 50 µL PCR reactions to add DNA homologous with the pCTCON vector to the ends of the amplified prey DNA. PCR products were combined and purified using phenol:chloroform extraction. DNA was concentrated using ethanol precipitation where a 1/10 volume of potassium acetate was added to the extracted DNA along with 10 µL of linear acrylamide followed by 2 volumes of ice-cold ethanol. The mixture was incubated at -20°C overnight followed by centrifugation at 15,000 g for 10 minutes the next day. After, the pellet was washed once with 70% ethanol followed by a 100% ethanol wash and allowed to dry prior to resuspension in water. PCR amplification and the DNA purification/concentration steps were repeated to obtain enough DNA for library construction.

In parallel, the NanoLuc protein was cloned into a dual display T2A construct that uses pCTCON as its backbone to generate pCTCON-T2A-NanoLuc. A similar procedure was used as described for the pCT302-T2A-NanoLuc construct. However, the pCTCON-T2A-SSoFe2-hFc construct as previously described in Bacon *et al*. was digested (4). 40 µg of pCTCON-T2A-NanoLuc was digested with NheI and BamHI and concentrated by phenol:chloroform extraction and ethanol precipitation.

The yeast-displayed cDNA library was generated by completing four electroporation reactions using a Bio-Rad Gene Pulser system (Bio-Rad) where 12 µg of amplified cDNA PCR product and 4 µg of digested vector was added to 400 µL of electrocompetent EBY100 cells. The electroporation settings used were 2.5 KV, 25 µF, and 250 Ω. An additional electroporation using only the digested vector was also performed. The number of transformed cells was estimated as $1x10^7$ as determined by plating serial dilutions of the combined transformation reactions onto SDCAA plates. However, the manufacturer of the cDNA library used for prey DNA amplification listed the diversity of their library as $6.64x10^6$.

*Plasmid construction for dual display of bait protein and SsoFe2 as yeast surface fusions*

The cDNA library screens were carried out against magnetic yeast co-displaying the bait protein and SsoFe2. To generate these cells, DNA encoding the bait proteins was cloned into the pCT302-SsoFe2-T2A-TOM22 plasmid between the NheI and BamHI sites as previously described (4). This plasmid confers the concurrent display of a target protein of interest and SsoFe2 as Aga2-yeast surface fusions using a T2A ribosomal skipping peptide. The bait proteins cloned into this construct include the WW domain of YAP (F164 – Q270) and SMAD3 by amplifying from gene block 3 using primers Pf8 and Pr8 and gene block 4 using Pf9 and Pr9, respectively. The WW domains of YAP were cloned using homologous recombination while SMAD3 was cloned via restriction digest using the NheI and BamHI sites. For homologous recombination, 200 ng of cut plasmid and 250 ng of insert were transformed into competent EBY100 cells generated using the Frozen-EZ Yeast Transformation Kit II. The plasmid and insert were concentrated using ethanol precipitation to ensure the total volume of DNA used in the transformation was less than 5 µL.

*Cloning to perform qYY2H assays on identified cDNA bait-prey pairs*

Some clones obtained from the cDNA screens were initially out of frame with the luciferase DNA due to the length of their poly A tail. For these constructs, the protein encoded by the cDNA was expressed correctly on the yeast surface, but luciferase was not displayed. Cloning was carried in a similar fashion as previously described using primers that changed the length of each clone's poly A tail. DNA encoding the in frame clones was inserted between the NheI and BamHI sites of pCTCON-T2A-NanoLuc.

All positive control peptides and proteins were cloned into the pCT302-T2A-NanoLuc construct via the NheI and BamHI sites as previously described. PTCH was amplified using gene block 5 with primers Pf10 and Pr10 while SMAD7 was amplified using gene block 6 with primers Pf11 and Pr11. The inserts were incorporated into the plasmid backbone using homologous recombination as previously described. The SMAD binding domain of SARA (A664 – C720) was amplified from gene block 7 using primers Pf12 and Pr12 and inserted into the plasmid backbone using restriction digest cloning via the NheI and BamHI sites.

*qYY2H assays to analyze identified cDNA bait-prey pairs*

A similar procedure as the other qYY2H assays was performed where $1x10^7$ magnetic bait cells (SMAD3 or WW domains of YAP) were incubated with $1x10^7$ prey cells co-expressing NanoLuc for 1 hour at room temperature along with $1x10^9$ EBY100. Complete titration curves were generated for prey cells expressing BRD7 and GGNBP2 binding to magnetic bait cells expressing SMAD3 as previously described. The individual titration curves from each repeat are described in **Figs. S18-S19** along with details on which points were included when fitting the monovalent binding isotherm to predict $K_{D,MV}$. **Equation S5** was used to estimate $K_D$ values for these prey.

*Cloning for analysis of bait-prey pairs that depend on post-translational modifications*

All prey proteins were cloned into pCTCON-T2A-NanoLuc construct via the NheI and BamHI sites as previously described. The SH2 domains of APS (E400 – T526) were amplified using gene block 8 with primers Pf13 and Pr13 (17). tSH2 was amplified using primers Pf14 and Pr14 while cSH2 and mSH2 were amplified using primers Pf15 and Pr14. SPY992 was amplified using primers Pf4 and Pr4. The DNA encoding tSH2, cSH2, mSH2, and SPY992 was amplified from plasmids described in Tiruthani *et al* (18).

The plasmid conferring expression of an EGFR peptide (F1163 – F1183, assuming no signal peptide) phosphorylated at the 1173 tyrosine was cloned in multiple steps. First, gene block 9 was amplified with primers Pf16 and Pr16. This DNA piece was inserted into the pCTCON plasmid via the AgeI and KpnI sites to generate pCTCON-AblTK. This part of the plasmid affords expression of the Abelson tyrosine kinase (AblTK) in the endoplasmic reticulum under the direction of the Gal10 promoter. Next, gene block 10 was amplified with primers Pf17 and Pr17 and inserted into the pCTCON-AblTK plasmid via the EcoRI and XhoI sites to generate pCTCON-AblTK-EGFR-T2A. Gene block 10 encodes the EGFR peptide substrate to be modified by AblTK as an Aga2 fusion as well as an Aga2-SsoFe2 fusion both under the direction of the Gal1 promoter. The DNA encoding the EGFR peptide substrate and SsoFe2 fusions are separated by a T2A ribosomal skipping peptide sequence that affords separate products of these two fusions after translation. N-terminal ER targeting tags are included for all expressed proteins as well as C-terminal ER retention tags to increase the residence time of the proteins within the ER. The final plasmid construct is described in **Fig. S7**.

In a similar manner, pCTCON-EGFR-T2A was generated by only inserting gene block 10 between the EcoRI and XhoI sites of pCTCON. This plasmid encodes a non-phosphorylated version of the described EGFR peptide as well as SsoFe2 as Aga2-yeast surface fusions.

*qYY2H assays for binding analysis of bait-prey pairs that depend on post-translational modifications*

Yeast cells containing the pCTCON-AblTK-EGFR-T2A plasmid were induced for protein expression for at least 24 hours at 20°C. These bait cells were magnetized as previously described. A similar procedure as the other qYY2H assays was performed where $1x10^7$ magnetic bait cells were incubated with $1x10^7$ prey cells co-expressing NanoLuc for 1 hour at room temperature along with $1x10^9$ EBY100.

A fraction of cells from the population that co-expresses Abelson kinase and the substrate peptide (pCTCON-AblTK-EGFR-T2A) will display non-phosphorylated peptides. Accordingly, flow cytometry analysis was used to estimate the percentage of peptide displaying cells that are express phosphorylated versions of the peptide. Peptide display, irrespective of phosphorylation state, was analyzed by labeling with an anti-c-myc antibody. Display of the phosphorylated peptide was analyzed by labeling with an anti-phosphotyrosine antibody. Briefly, $2x10^6$ cells were labeled with a 1:100 dilution of chicken anti-c-myc antibody (Thermo Fisher Scientific) or a 1:50 dilution of Alexa Fluor 647 anti-phosphotyrosine (Biolegend) for 15 minutes at room temperature. Subsequently, cells were washed and secondary labeling was carried out for the cells labeled with the anti-c-myc antibody using a 1:250 dilution of goat-anti-chicken 488 (Immunoreagents) for 10 minutes on ice. All labeling were conducted in 50 μL of 0.1% PBSA, and washes with 0.1% PBSA took place after each labeling. Cells were analyzed using a Miltenyi Biotec MACsQuant VYB cytometer. The cells were not double labeled for c-myc and phosphorylation expression as it was observed that the binding of the anti-phosphotyrosine antibody decreased during double labeling due to steric hindrance. Similar analysis was completed for cells containing the pCTCON-EGFR-T2A plasmid (no modifying enzyme) to quantify their c-myc expression. For the pCTCON-AblTK-EGFR-T2A cells, 75.2% of the total cell population expressed a peptide while only 37.6% of the total cell population expressed a phosphorylated version of the peptide **(Figs. 5A-5B)**. Therefore, only around ~50% of the peptide-displaying pCTCON-AblTK-EGFR-T2A cells are actually displaying a phosphorylated version of the peptide.

Prey cell recovery by the phosphorylated bait cell population was normalized by the percentage of peptide-displaying cells expressing a phosphorylated version of the peptide. This normalization took place as it is likely a higher number of prey cells could have been recovered if all cells within the bait population displayed a phosphorylated version of the EGFR peptide.

**Supplementary Figures and Tables**

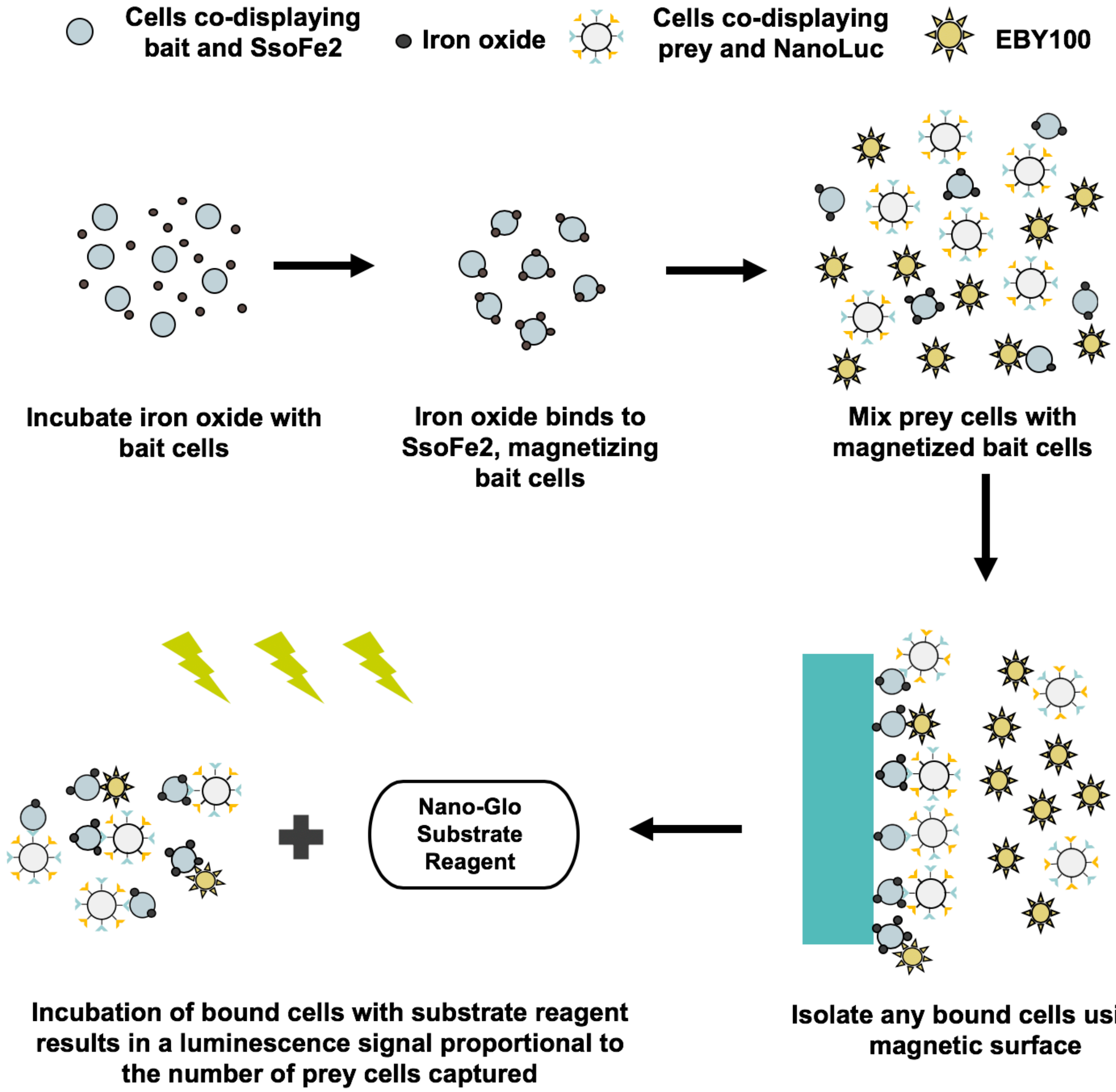

**Figure S1**. Detailed strategy for quantitative yeast-yeast two hybrid assay. The steps include the magnetization of the bait cells after incubation with iron oxide, incubation of the bait and prey cells, isolation of any prey cells bound by the magnetic bait cells using a magnet, and quantification of prey cell capture using the luminescence read out after addition of the Nano-Glo reagent.

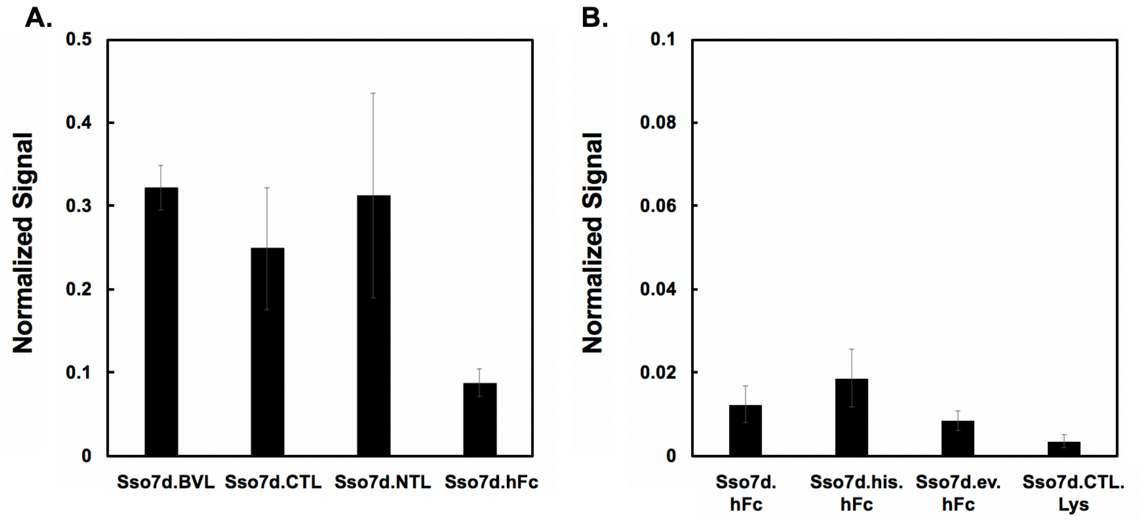

**Figure S2**. Evaluation of GOx as a reporter enzyme for the quantitative assessment of bait-prey interactions as a function of prey cell recovery. Prey proteins were expressed as yeast surface fusions along with GOx. **(A)** Normalized GOx signal proportional to recovery of prey cells by magnetic beads functionalized with lysozyme. The prey cells displayed either Sso7d.BVL.Lys, Sso7d.CTL.Lys, Sso7d.NTL.Lys, or an irrelevant protein, Sso7d.hFc. **(B)** Normalized GOx signal proportional to recover of prey cells by magnetic yeast displaying the bait protein (hFc). The prey cells displayed either Ssso7d.hFc, Sso7d.his.hFc, Ssso7d.ev.hFc, or an irrelevant protein, Sso7d.CTL.Lys. GOx signals were normalized throughout by the mean fluorescence associated with prey expression (detected using an anti-c-myc antibody) and by a GOx activity control. Error bars represent the standard error of the mean for three repeats throughout.

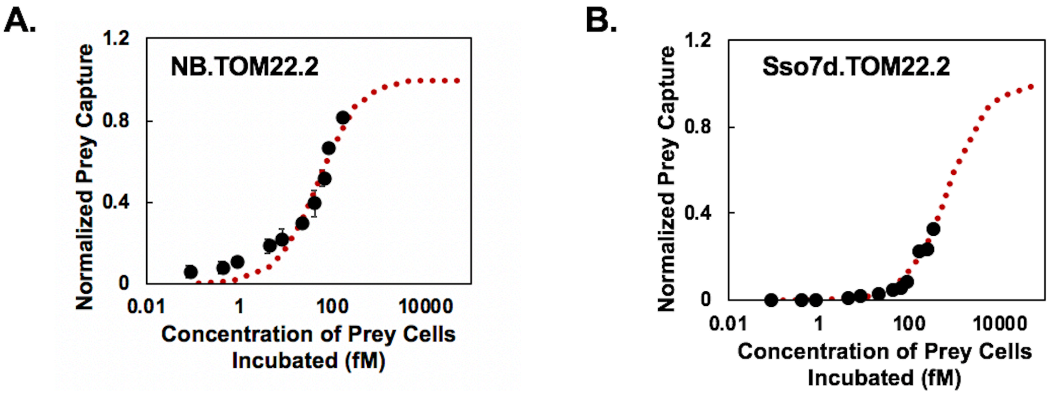

**Figure S3**. Titration curves describing the binding of prey cells expressing **(A)** NB.TOM22.2 or **(B)** Sso7d.TOM22.2 as a function of the concentration of prey cells incubated were generated for a given concentration of magnetic bait cells expressing TOM22. A global fit was used to predict $K_{D,MV}$ using the proposed non-linear, multivalent binding model as represented by the red line. Error bars represent the standard error of the mean for 3 or 4 repeats.

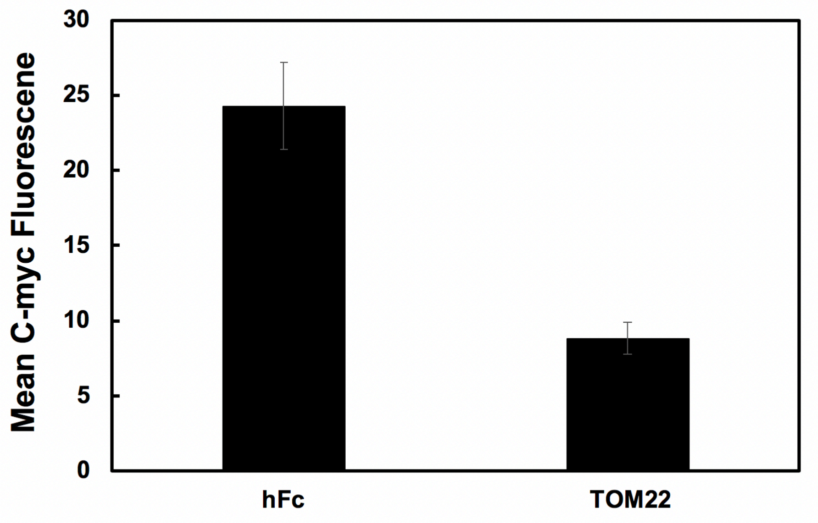

**Figure S4**. Comparison of bait expression levels between cells co-expressing either hFc and SsoFe2 or TOM22 and SsoFe2. The expression of the bait protein was quantified by labeling with an anti-c-myc antibody followed by detection with a goat-anti-chicken secondary antibody and flow cytometry analysis. The expression of the hFc bait on the surface of yeast is ~2.7 fold higher than the expression of the TOM22 bait on the yeast surface. Error bars represent the standard error of the mean for eight independent replicates.

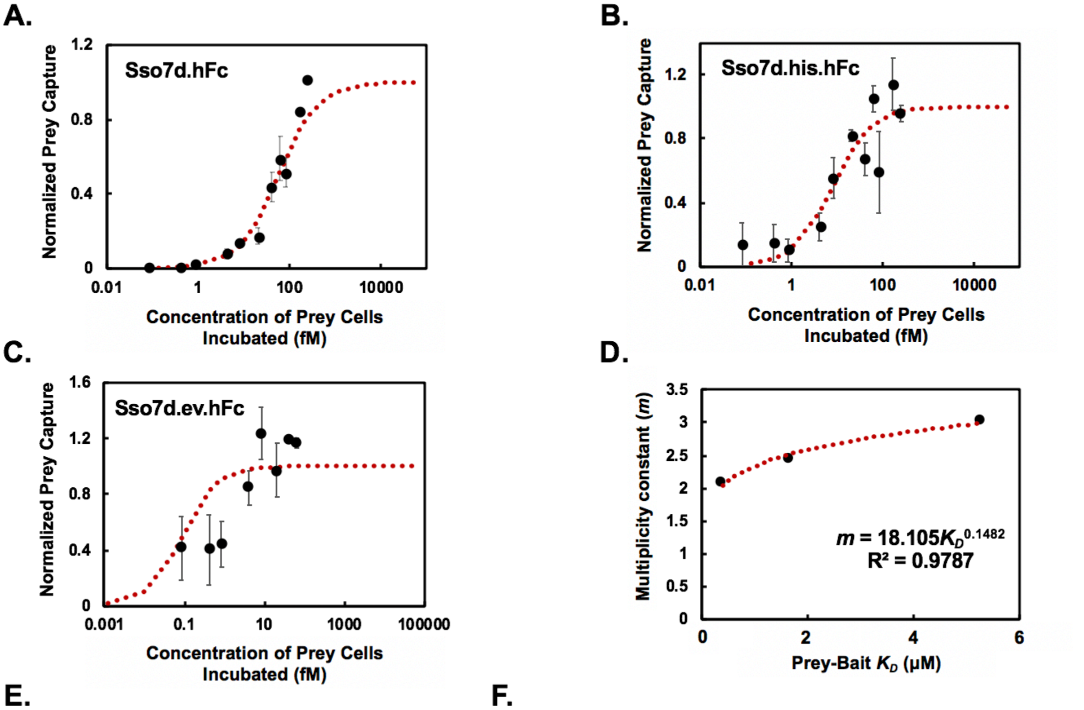

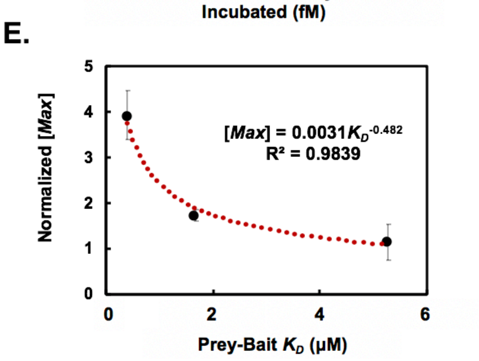

**F.**

| | $K_{D,MV}$ (fM) | YST $K_D$ (nM) | $m$ |
|---|---|---|---|
| **Sso7d.hFc** | 58.4 (47.5 – 72.3) | 400 (366 – 437) | 2.07 |
| **Sso7d.his.hFc** | 8.1 (3.9 – 16.1) | 1670 (1337 – 3000) | 2.44 |
| **Sso7d.ev.hFc** | 0.087 (0.013 – 0.34) | 5280 (4500 – 6700) | 3.04 |

**Figure S5**. A semi-empirical framework enables the development of a relationship between monovalent $K_D$ and multivalent $K_{D,MV}$ when the bait protein is immobilized on magnetic beads. Titration curves describing the binding of prey cells as a function of the concentration of prey cells incubated were generated for a given concentration of magnetic beads immobilized with the bait protein (biotinylated hFc). For each repeat, the concentration of prey cells captured is normalized by the estimated maximum concentration of prey cells captured, [*Max*]. Titration curves are shown for **(A)** Sso7d.hFc, **(B)** Sso7d.his.hFc, and **(C)** Sso7d.ev.hFc. **(D)** A global non-linear regression was used to estimate $K_{D,MV}$, the equilibrium dissociation constant describing multivalent, cell-magnetic bead binding, as well as the multiplicity constant $m$. A plot of $m$ vs. $K_D$ is shown and fits a power law model. $K_D$ values for these bait-prey interactions had been previously estimated from soluble yeast surface titration (YST $K_D$). **(E)** A plot of normalized [*Max*] values vs. $K_D$ fits a power law model. [*Max*] values for each repeat were normalized by the average number of prey cells captured across each concentration of prey cells incubated. **(F)** Experimentally determined $K_{D,MV}$ and calculated values of $m$ based on previous estimates of $K_D$ (YST $K_D$) are shown . Values in parentheses represent the bounds of a 68% confidence interval. Error bars represent the standard error of the mean for three repeats for all data shown.

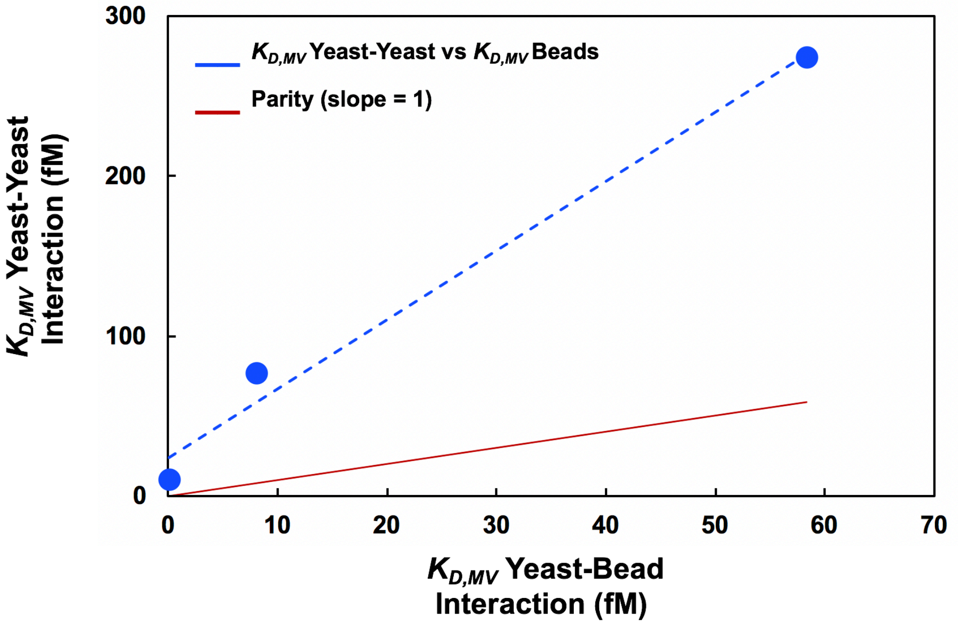

**Figure S6.** Comparison of $K_{D,MV}$ for yeast-yeast interactions (y-axis) and $K_{D,MV}$ for yeast-bead interactions (x-axis). The interaction of prey cells displaying Sso7d.hFc, Sso7d.his.hFc, or Sso7d.ev.hFc with either magnetic cells expressing the hFc bait or magnetic beads functionalized with soluble hFc were considered (blue). Included in the figure is a parity line (red) that details the trend if the $K_{D,MV}$ for the yeast-yeast interaction was the same as the $K_{D,MV}$ for yeast-bead interaction. Accordingly, the $K_{D,MV}$ is significantly lower for yeast-bead interaction relative to yeast-yeast interaction.

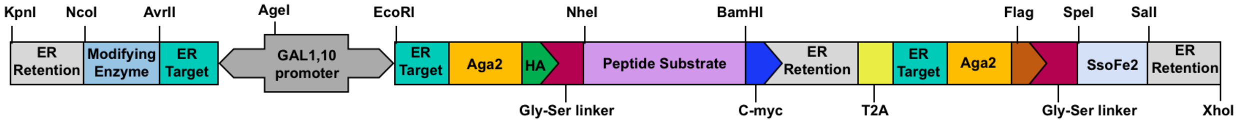

**Figure S7**. Plasmid design for co-expression of an enzyme modified bait peptide and SsoFe2, an iron oxide binding protein, as fusions to Aga2 for display on the yeast surface. The DNA encoding the peptide substrate and SsoFe2 is under the control of the Gal1 promoter and is separated by a T2A ribosomal skipping peptide, which affords separate translation of the fusion proteins. A bait-modifying enzyme is encoded downstream of the Gal10 promoter. Each protein is trafficked to the ER by an included N-terminal targeting sequence as well being retained in the ER for a longer than average residence time due to an included C-terminal ER retention sequence. This increases the probability that the modifying enzyme will have time to interact with the peptide substrate. Ultimately, the modified peptide substrates along with SsoFe2 are trafficked to the yeast cell surface for display.

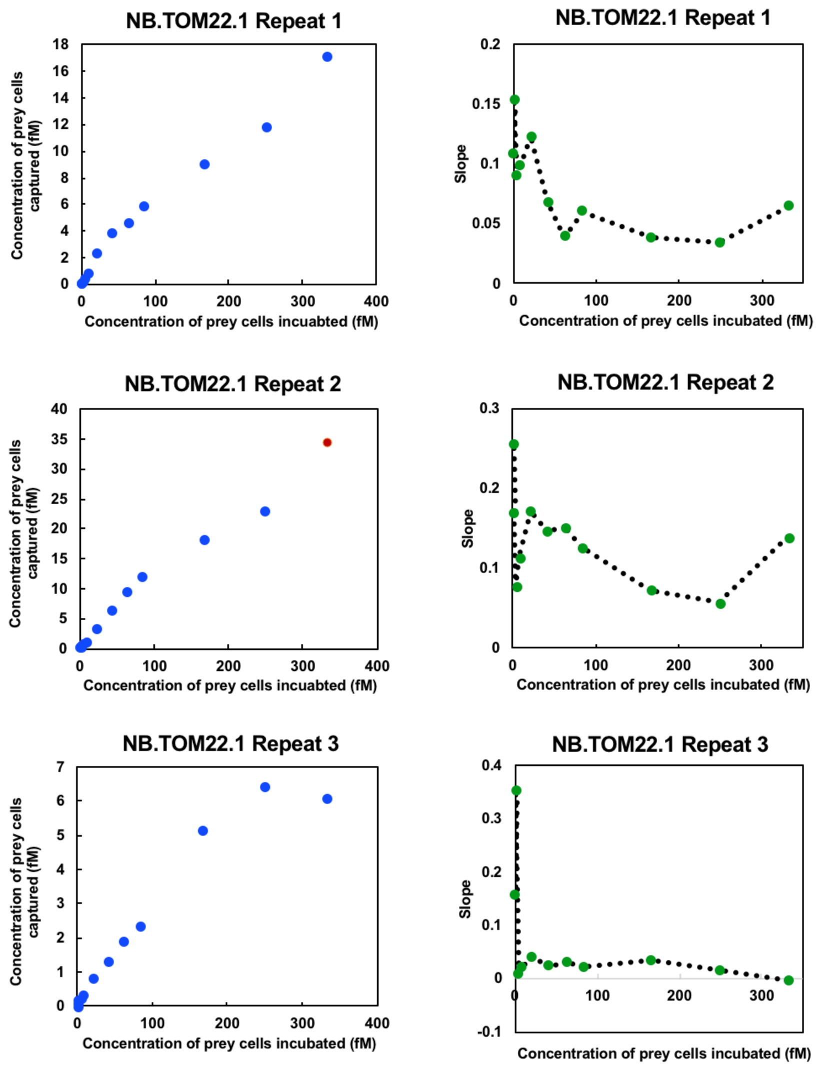

**Figure S8.** Titration curves (left) generated for a NB.TOM22.1 prey comparing the concentration of prey cells captured as a function of the concentration of prey cells incubated with magnetic yeast cells expressing the bait, TOM22. The number of magnetic bait cells displaying TOM22 was held constant. A plot of the slope between adjacent titration curve data points was also generated (right). The slope plot was used to determine which points to include when applying the monovalent binding isotherm to the titration curve data. Typically, points were ignored after saturation was met (slope ~0). Non-specific binding of the prey cells can occur after saturation is met. For the titration curve plots, blue points were included when applying the monovalent binding isotherm while red points were discarded.

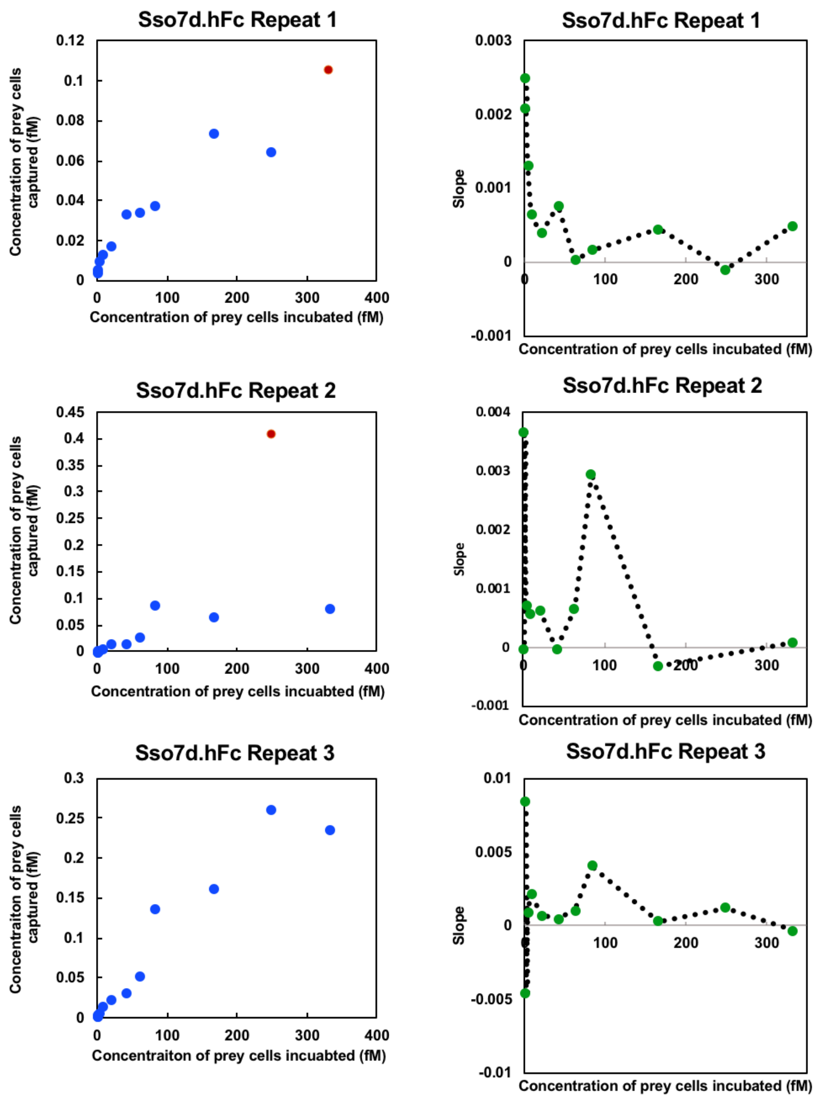

**Figure S9.** Titration curves (left) generated for a Sso7d.hFc prey comparing the concentration of prey cells captured as a function of the concentration of prey cells incubated with magnetic yeast cells expressing the bait, hFc. The number of magnetic bait cells displaying hFc was held constant. A plot of the slope between adjacent titration curve data points was also generated (right). The slope plot was used to determine which points to include when applying the monovalent binding isotherm to the titration curve data. Typically, points were ignored after saturation was met (slope ~0). Non-specific binding of the prey cells can occur after saturation is met. For the titration curve plots, blue points were included when applying the monovalent binding isotherm while red points were discarded.

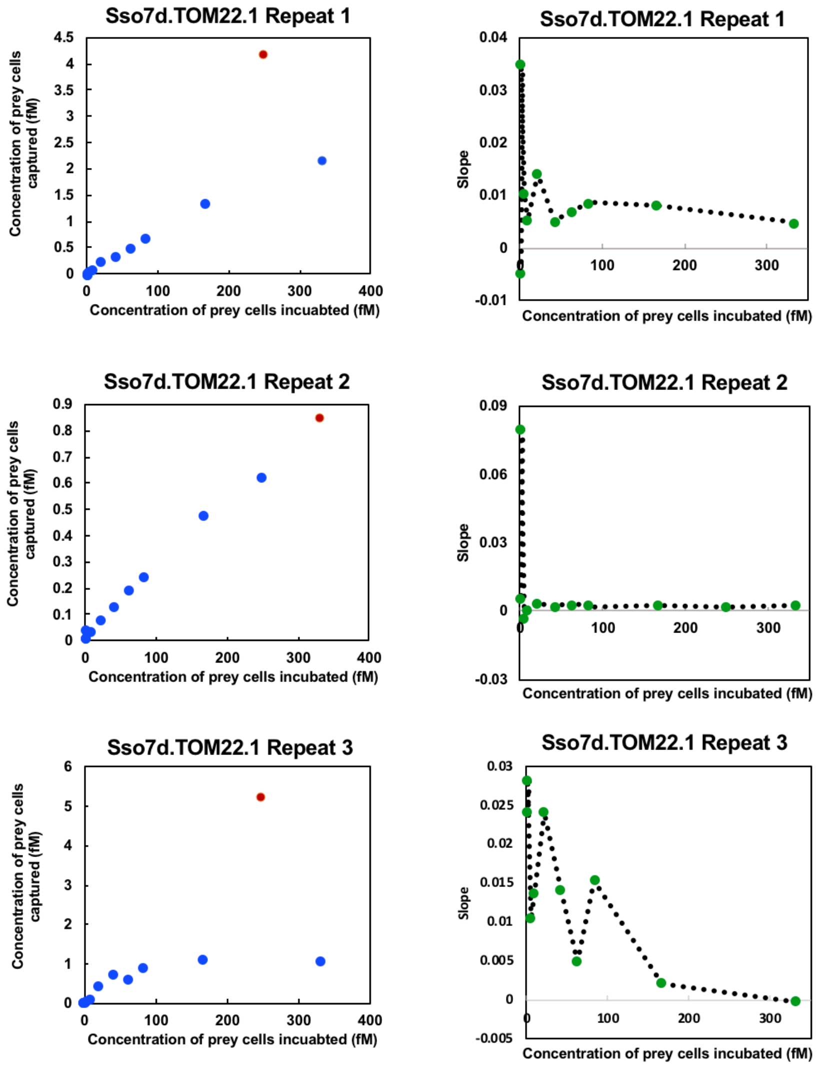

**Figure S10.** Titration curves (left) generated for a Sso7d.TOM22.1 prey comparing the concentration of prey cells captured as a function of the concentration of prey cells incubated with magnetic yeast cells expressing the bait, TOM22. The number of magnetic bait cells displaying TOM22 was held constant. A plot of the slope between adjacent titration curve data points was also generated (right). The slope plot was used to determine which points to include when applying the monovalent binding isotherm to the titration curve data. Typically, points were ignored after saturation was met (slope ~0). Non-specific binding of the prey cells can occur after saturation is met. For the titration curve plots, blue points were included when applying the monovalent binding isotherm while red points were discarded.

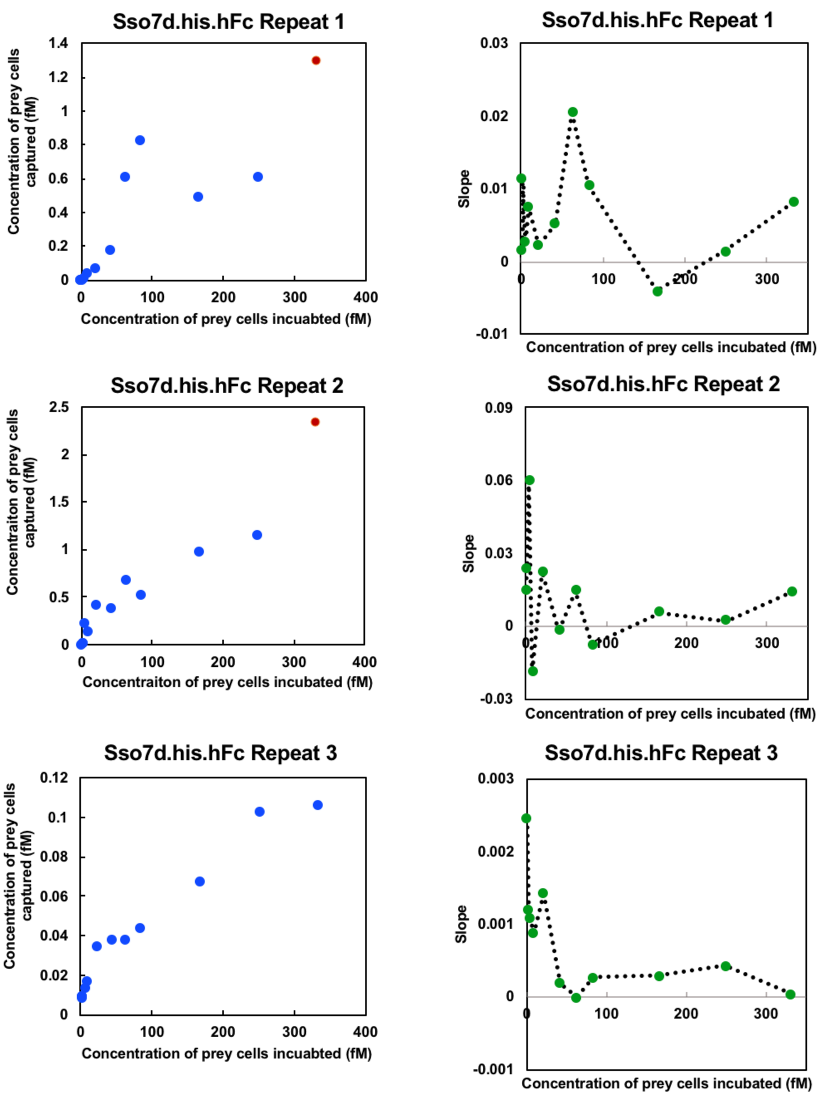

**Figure S11.** Titration curves (left) generated for a Sso7d.his.hFc prey comparing the concentration of prey cells captured as a function of the concentration of prey cells incubated with magnetic yeast cells expressing the bait, hFc. The number of magnetic bait cells displaying hFc was held constant. A plot of the slope between adjacent titration curve data points was also generated (right). The slope plot was used to determine which points to include when applying the monovalent binding isotherm to the titration curve data. Typically, points were ignored after saturation was met (slope ~0). Non-specific binding of the prey cells can occur after saturation is met. For the titration curve plots, blue points were included when applying the monovalent binding isotherm while red points were discarded.

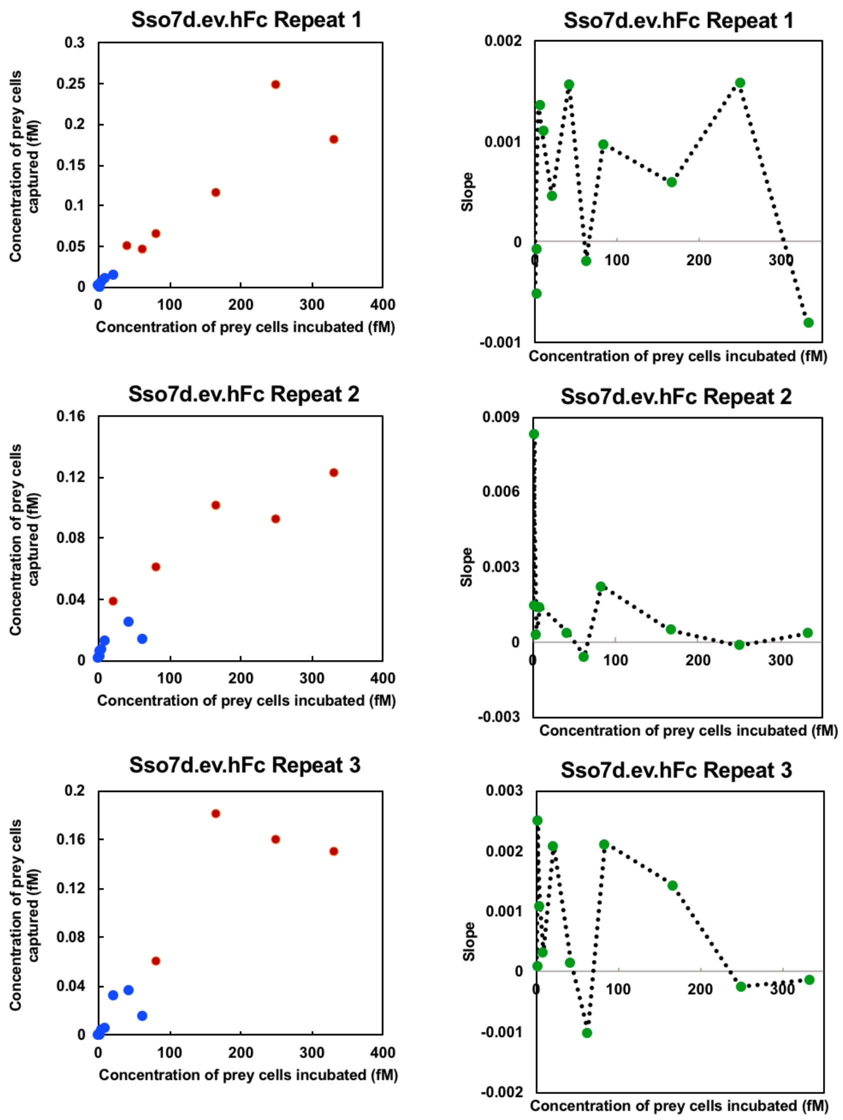

**Figure S12.** Titration curves (left) generated for a Sso7d.ev.hFc prey comparing the concentration of prey cells captured as a function of the concentration of prey cells incubated with magnetic yeast cells expressing the bait, hFc. The number of magnetic bait cells displaying hFc was held constant. A plot of the slope between adjacent titration curve data points was also generated (right). The slope plot was used to determine which points to include when applying the monovalent binding isotherm to the titration curve data. Typically, points were ignored after saturation was met (slope ~0). Non-specific binding of the prey cells can occur after saturation is met. For the titration curve plots, blue points were included when applying the monovalent binding isotherm while red points were discarded.

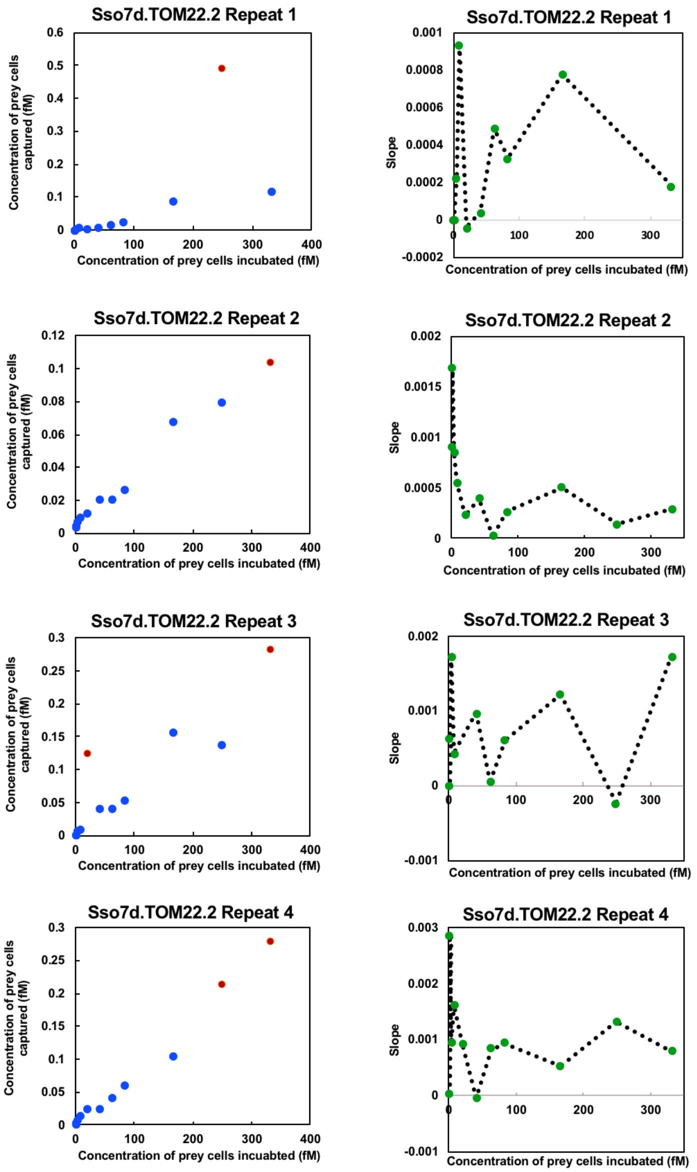

**Figure S13.** Titration curves (left) generated for a Sso7d.TOM22.2 prey comparing the concentration of prey cells captured as a function of the concentration of prey cells incubated with magnetic yeast cells expressing the bait, TOM22. The number of magnetic bait cells displaying TOM22 was held constant. A plot of the slope between adjacent titration curve data points was also generated (right). The slope plot was used to determine which points to include when applying the monovalent binding isotherm to the titration curve data. Typically, points were ignored after saturation was met (slope ~0). Non-specific binding of the prey cells can occur after saturation is met. For the titration curve plots, blue points were included when applying the monovalent binding isotherm while red points were discarded.

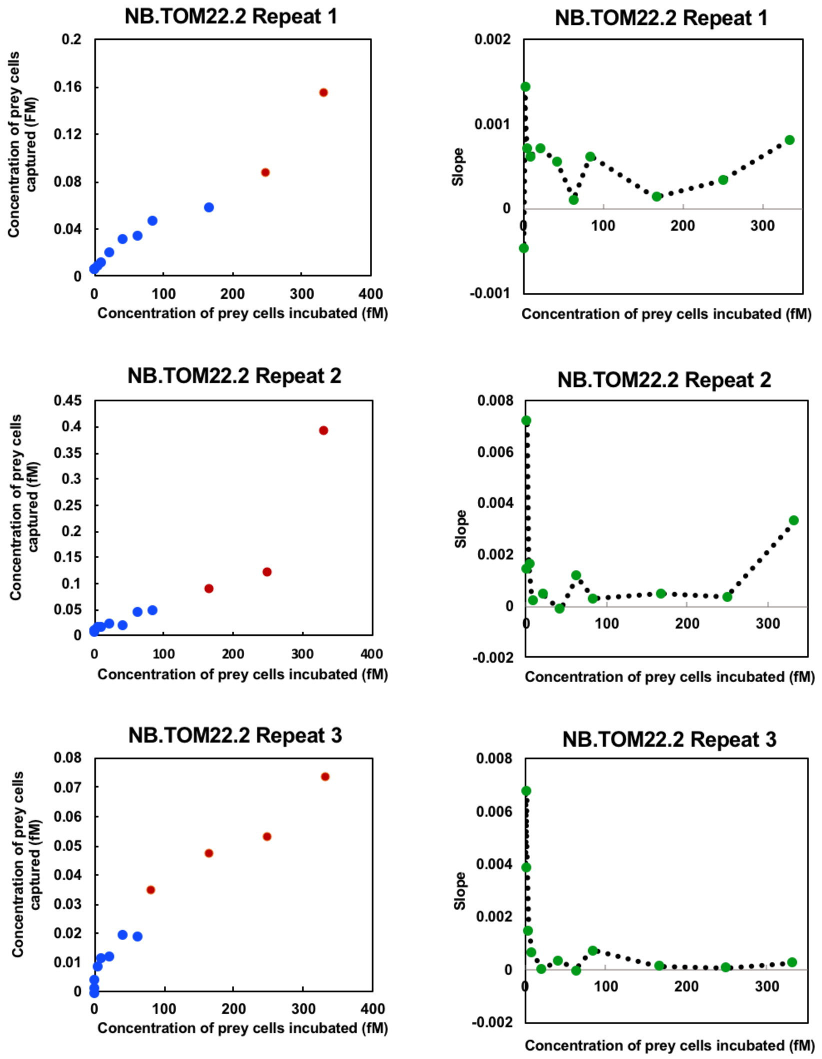

**Figure S14.** Titration curves (left) generated for a NB.TOM22.2 prey comparing the concentration of prey cells captured as a function of the concentration of prey cells incubated with magnetic yeast cells expressing the bait, TOM22. The number of magnetic bait cells displaying TOM22 was held constant. A plot of the slope between adjacent titration curve data points was also generated (right). The slope plot was used to determine which points to include when applying the monovalent binding isotherm to the titration curve data. Typically, points were ignored after saturation was met (slope ~0). Non-specific binding of the prey cells can occur after saturation is met. For the titration curve plots, blue points were included when applying the monovalent binding isotherm while red points were discarded.

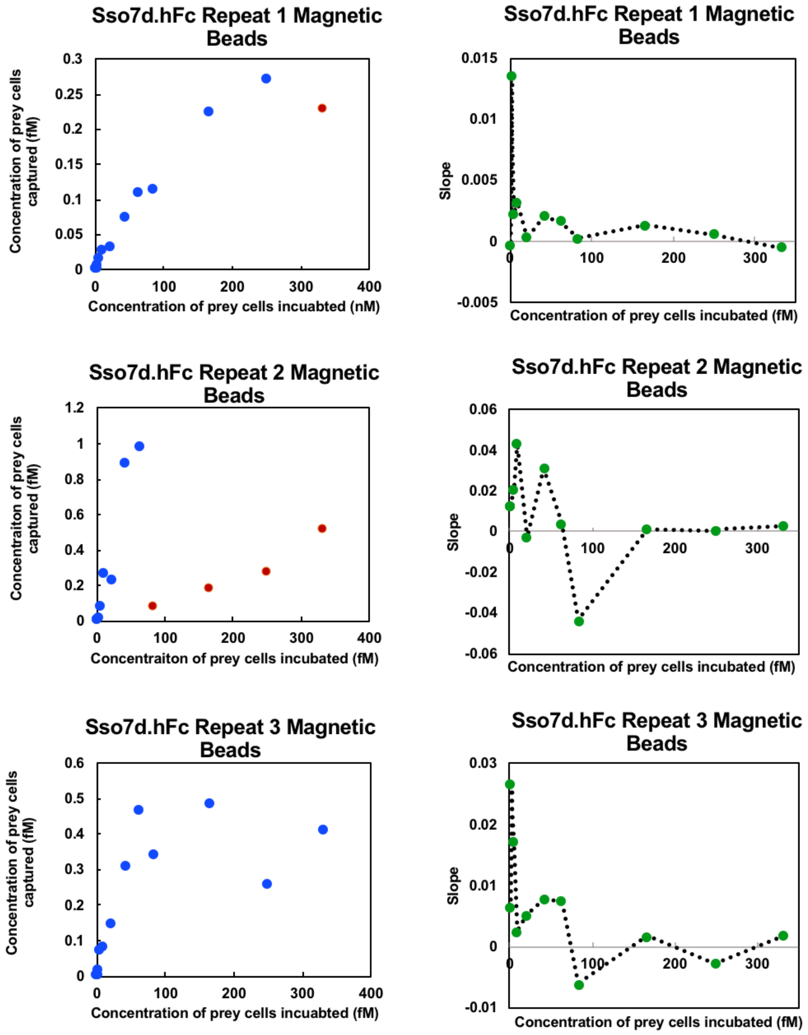

**Figure S15.** Titration curves (left) generated for a Sso7d.hFc prey comparing the concentration of prey cells captured as a function of the concentration of prey cells incubated with magnetic beads functionalized with the bait, hFc. The number of magnetic beads functionalized with hFc was held constant. A plot of the slope between adjacent titration curve data points was also generated (right). The slope plot was used to determine which points to include when applying the monovalent binding isotherm to the titration curve data. Typically, points were ignored after saturation was met (slope ~0). Non-specific binding of the prey cells can occur after saturation is met. For the titration curve plots, blue points were included when applying the monovalent binding isotherm while red points were discarded.

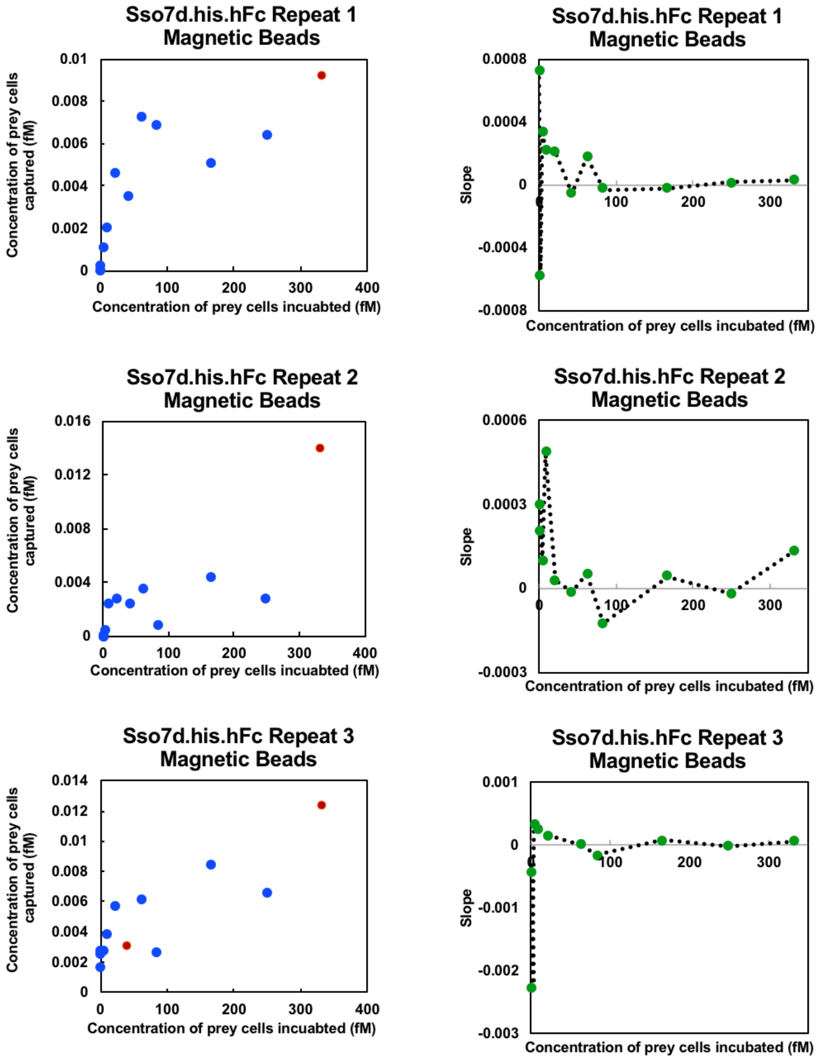

**Figure S16.** Titration curves (left) generated for a Sso7d.his.hFc prey comparing the concentration of prey cells captured as a function of the concentration of prey cells incubated with magnetic beads functionalized with the bait, hFc. The number of magnetic beads functionalized with hFc was held constant. A plot of the slope between adjacent titration curve data points was also generated (right). The slope plot was used to determine which points to include when applying the monovalent binding isotherm to the titration curve data. Typically, points were ignored after saturation was met (slope ~0). Non-specific binding of the prey cells can occur after saturation is met. For the titration curve plots, blue points were included when applying the monovalent binding isotherm while red points were discarded.

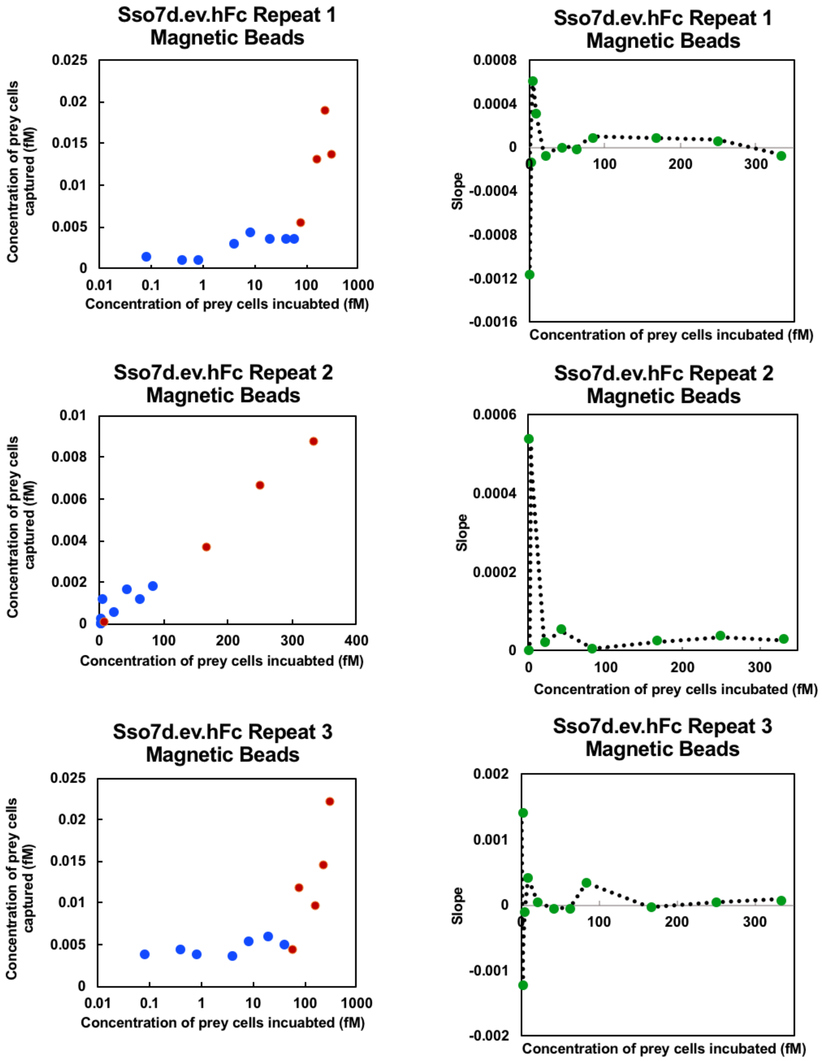

**Figure S17.** Titration curves (left) generated for a Sso7d.ev.hFc prey comparing the concentration of prey cells captured as a function of the concentration of prey cells incubated with magnetic beads functionalized with the bait, hFc. The number of magnetic beads functionalized with hFc was held constant. A plot of the slope between adjacent titration curve data points was also generated (right). The slope plot was used to determine which points to include when applying the monovalent binding isotherm to the titration curve data. Typically, points were ignored after saturation was met (slope ~0). Non-specific binding of the prey cells can occur after saturation is met. For the titration curve plots, blue points were included when applying the monovalent binding isotherm while red points were discarded.

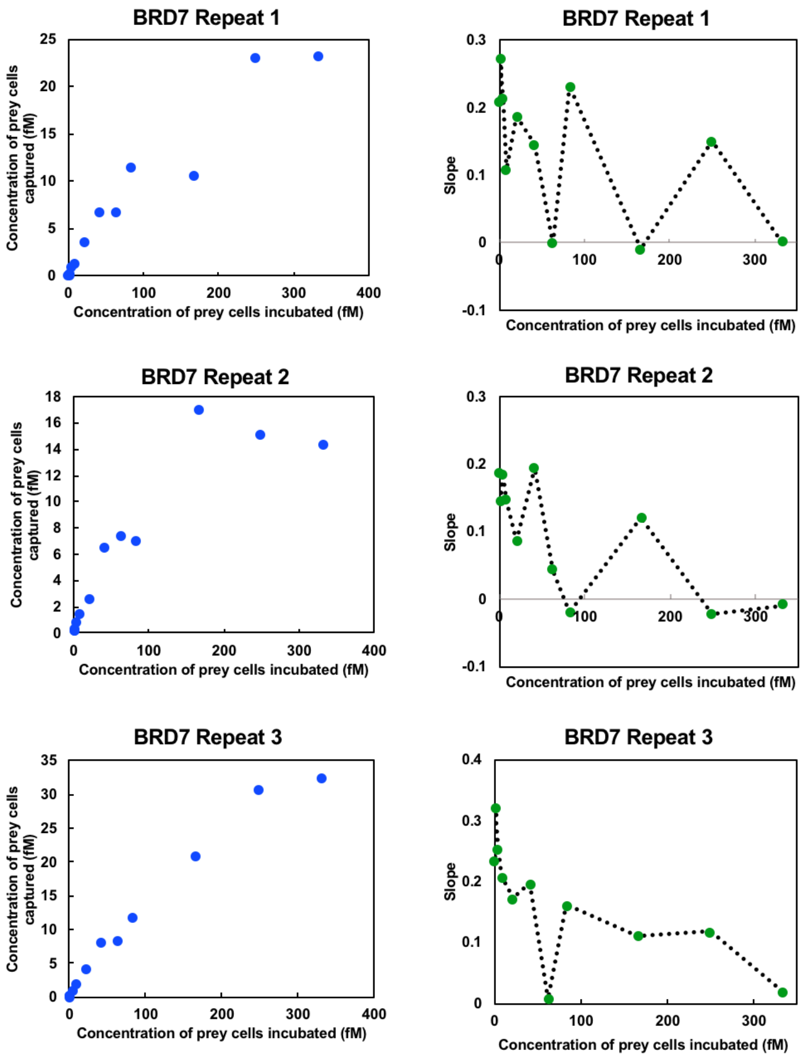

**Figure S18.** Titration curves (left) generated for a BRD7 prey comparing the concentration of prey cells captured as a function of the concentration of prey cells incubated with magnetic yeast cells expressing the bait, SMAD3. The number of magnetic bait cells displaying SMAD3 was held constant. A plot of the slope between adjacent titration curve data points was also generated (right). The slope plot was used to determine which points to include when applying the monovalent binding isotherm to the titration curve data. Typically, points were ignored after saturation was met (slope ~0). Non-specific binding of the prey cells can occur after saturation is met. For the titration curve plots, blue points were included when applying the monovalent binding isotherm while red points were discarded.

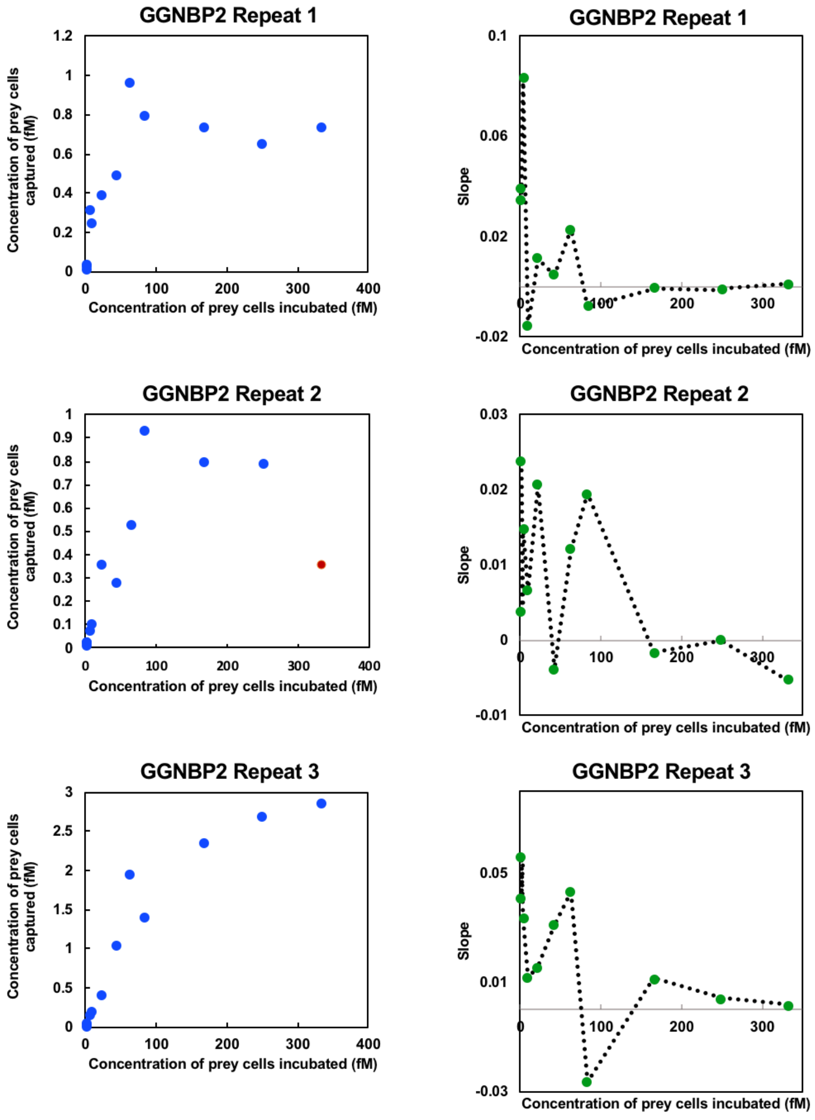

**Figure S19.** Titration curves (left) generated for a GGNBP2 prey comparing the concentration of prey cells captured as a function of the concentration of prey cells incubated with magnetic yeast cells expressing the bait, SMAD3. The number of magnetic bait cells displaying SMAD3 was held constant. A plot of the slope between adjacent titration curve data points was also generated (right). The slope plot was used to determine which points to include when applying the monovalent binding isotherm to the titration curve data. Typically, points were ignored after saturation was met (slope ~0). Non-specific binding of the prey cells can occur after saturation is met. For the titration curve plots, blue points were included when applying the monovalent binding isotherm while red points were discarded.

**Table S1**. Putative interaction partners for SMAD3 isolated from a yeast-displayed cDNA library screened against magnetic yeast cells displaying SMAD3. Prey cells displaying the protein encoded by the uncut vector used for library construction were isolated due to nonspecific binding.

| ***Protein*** | ***Abbreviation*** | ***Occurrence*** | ***Protein Subunit Amino Acid Sequence*** |
|---|---|---|---|
| NIPBL cohesin loading factor | NIPBL | 1 | ETPKQKSEGRPETPKQKGDGRPETPKQKSEGRPETPKQKGEGRPETPKHRHENRRDSGKPSTEKKPDVSKHKQDIKSDSPRLKSERAEALKQRPDGRWESLRRDHDSKQKSDDRGESERHRGDQSRVRRPETLRSSSRNDHSTKSDAGKTEK |
| Bromodomain containing protein 7 | BRD7 | 1 | GGPVGHGQEAQEAQVGPPLLRGVRGEALKLVLKVGGSEVTELSTGSSGHDSSLFEDRSDHDKHKDRKRKKRKKGEKQAPGEEKGRKRRRVKEDKKKRDRDRAENEVDRDLQCHVPIRLDLPPEKPLTSSLAKQEEVEQTPLQEALNQLMRQLQRKDPSAFFSFPVTDFIAPGYSMIIKHPMDFSTMKEKIKNNDYQSIEELKDNFKLMCTNAMIYNKPETIYYKAAKKLLHSGMKILSQERIQSLKQSIDFMSDLQKTRKQKERTDACQSGEDSGCWQREREDSGDAGKTEKK |
| Gametogenetin binding protein 2 | GGNBP2 | 1 | EQLCEEFSEEERVRELKQEKKRQKRKNRK |

| | | | |
|---|---|---|---|
| Endothelin 1 | EDN1 | 1 | NTPERVVPYGLGGSSRSKRSLKDLLPNKATDQAVRCQCAHQKDKKCWKNRK |
| Mitochondrial calcium uniporter | MCU | 1 | RHHVLSQLHHPRCWKNRKKD |
| Dead box polypeptide 18 | DDX18 | 1 | RHKPVPTSKWRCWKNRK |
| Pre-mRNA processing factor 40A | PRPF40A | 1 | AEWHYGRGKKKSKKRRHKSDSPESDAGKTEK |
| TATA-Box binding protein associated factor 2 | TAF2 | 1 | KKKKHKHKHKHKRKHDSKDKDREPFAFSSPASGRPIQMLEKQKK |
| Nucleotide binding protein 1 | NUBP1 | 1 | RVGRPRVQTPGQWGTGGTIATRCWKNRK |
| Uncut Vector | | 21 | |

**Table S2**. Putative interaction partners for the WW domains of YAP isolated from a yeast-displayed cDNA library screened against magnetic yeast displaying the bait protein. Some of the isolated prey contain previously identified binding motifs for the WW domain. Prey cells displaying the protein encoded by the uncut vector used for library construction were isolated due to nonspecific binding.

| Protein | Abbreviation | Occurrence | Binding Motif | Protein Subunit Amino Acid Sequence |
|---|---|---|---|---|
| Tata-Box binding protein associated factor 2 | TRF2 | 2 | RP | HKHKHKHKHKHDSKD KDREPFAFSSPASGRPI QMLEKQKK |
| Cyclin L2 | CCNL2 | 1 | PPR | AEGHYGRGGGSKSQS RSRSRSDSPPRQVHRG APYKGSEVRGSRKSKD CKYLTQKPHKSRSRSS SRSRSRSRERTDNAGK TEK |
| Nuclear speckle splicing regulatory protein 1 | NSRP1 | 1 | RPP, RP | AEWPLRPERSSSEERG LGTKHHSRGSQSRGHE HQDRQSRDQESCHKD RSHREEKSSHRHREAS HKDHYWKKHEQEDKL KGREQEERQDREGKR EKYSSREQERDRQRND HDRYSEKEKKRKEKEE HTKARRERCEDSGKH REREKPEGHGQSSERH RDRRESSPRSRPKDDL DQERSSKARNTEKDK GEQGKPSHSETSLATK |

| | | | | |
|---|---|---|---|---|
| | | | | HRLAEERPEKGSEQERPPEAVSKFAKRSNEETVMSARDRYLARQMARINAKTYIEKEDD |
| Ribosome protein L3 | RPL3 | 1 | None | VFAEHISDECKRRFYKNWHKSKKKAFTKYCKKWQDDTGKKQLEKDFNSMKKYCQVIRIIAHTQMLEKQKK |
| Chromatin target of PRMT1 | CHTOP | 1 | None | SHHIYKTTLRKHEKKSDRCWKNRK |
| Chromosome 12 RP23-33124 | | 1 | RP | RKCGRPAHSWKNRK |
| Chromosome 7, RP24-233E11 | | 1 | RP | EKKKKKKKPAHTHKHTITHISVYSNSATLLTRIN |
| Guanine nucleotide binding protein-like 2 | GNL2 | 1 | None | KEEKKTSAEDSDAAPTKKARKWDAQMEEEPSNKTQRMLTCKERGRAARQQQSKKVGVRYYETHNLKNRNRNKKKTSDSEGQKHRRNKFRQKQ |
| ADP-ribosylation factor like GTPAse 9 | ARL9 | 2 | None | KQGEGEREGEGRGEKEVCGTEKQEGERDRERRREREGGRSENQPRERERREEKEECGPGNQREWERQGEGGREKEECGPKNQGEWERQGEGGR |

| | | | | |
|---|---|---|---|---|
| | | | | EKEECGPETKENGKDK GKEEEKKKSVDLKTKE NGKDKGKEEGKKKSV DLKTKENEKDKGKEE EKKKSVDLKTKENGK DKGKEEEKKKSVDPK TKENEKDKGKEEEKK KSVDLKTKENGKDKG KEEEKKKSVDLKTKEN GKDKEKEEKKRKSAD LKTKETGKEKGRREK K |
| Transcription elongation regulator 1 | TCERG1 | 1 | PXY | REKKNKIMQAKEDFK KMMEEAKFNPRATFSE FAAKRAKDSRFKAIEK MKDREALFNEFVAAA RKKEKEDSKTRGEKIK SDFFELLSNHHLDSQS RWSKVKDKVESDPRY KAVDSSSMREDLFKQ YIEKIAKNLDSEKEKEL ERQARIEASLRERERE VQKARSEQTKEIDRER EQHKREEAIQNFKALL SDMVRSSDVSWSDTR RTLRKDHRWESGSLLE REEKEKLFNEHIEALT KKKKKKAASASRGWA SIRDPSSSSCR |
| Undigested vector | | 14 | | |

| | | |
|---|---|---|
| Small cDNA with no similarity | | 4 |

**Table S3**. List of gene fragments

| | |
|---|---|
| Gene Block 1 | CAACTTACCCCTACCTTCTACGACAATTCATGTCCTAACGTCTCAAACATAGTACGGGACACTATTGTCAATGAGTTACGATCGGACCCTAGAATCGCCGCGAGCATCCTTCGTCTTCACTTCCACGACTGCTTTGTTAACGGTTGTGACGCATCGATCTTGTTAGACAACACAACATCATTTCGAACAGAGAAAGATGCGTTTGGGAACGCAAACTCGGCGCGCGGATTTCCTGTGATTGACAGAATGAAGGCGGCCGTGGAGAGTGCATGCCCAAGAACTGTATCCTGCGCAGATCTGCTTACCATTGCAGCTCAACAATCTGTTACTTTGGCAGGAGGTCCCTCTTGGAGGGTTCCTTTGGGACGTCGAGACAGCCTACAAGCATTTTTAGATCTCGCGAATGCGAATCTTCCAGCTCCATTCTTCACACTTCCACAACTTAAGGATTCTTTTAGAAATGTTGGTTTAAACCGTTCTTCTGATCTCGTTGCCCTCAGCGGTGGTCACACATTTGGTAAAAATCAATGTCGATTTATTATGGACAGATTATACAACTTCAGCAACACCGGGTTACCCGACCCTACCCTCAACACTACTTACCTTCAAACTCTTCGTGGACTATGTCCCCTTAATGGCAACCTAAGCGCCTTGGTCGATTTCGATCTGCGTACGCCAACAATTTTCGATAACAAATACTATGTGAATCTTGAAGAGCAAAAAGGTCTCATCCAGAGTGATCAAGAGTTGTTCTCTAGCCCCAATGCCACTGACACAATCCCACTGGTGAGATCATTTGCTAATAGCACACAAACATTCTTCAACGCGTTTGTGGAGGCCATGGATAGGATGGGAAACATTACACCTCTTACAGGAACTCAAGGACAAATCAGGTTGAACTGTAGGGTGGTGAACTCCAACTCT |
| Gene Block 2 | ATGGTTTTCACCTTAGAGGATTTCGTAGGCGACTGGAGACAGACGGCAGGCTATAACTTGGACCAAGTGCTGGAACAGGGCGGTGTCAGCTCCTTATTCCAGAATCTGGGGGTATCCGTTACTCCTATACAGAGAATAGTTCTTTCCGGAGAAAACGGGCTAAAGATAGATATTCATGTCATTATTCCTTACGAAGGATTGTCCGGCGACCAAATGGGTCAAATCGAAAAGATTTTCAAGGTGGTTTACCCGGTAGACGATCATCACTTCAAGGTGATTTTGCATTACGGTACATTAGTTATTGACGGGGTAACGCCAAACATGATAGACTACTTTGGCAGACCCTACGAGGGAATTGCAGTTTTCGATGGTAAAAAGATCACGGTTACGGGTACTCTGTGGAA |

| | |
|---|---|
| | CGGGAATAAAATTATAGATGAGCGTTTAATCAATCCAGATGGATCACTACTTTTTAGAGTGACAATTAACGGTGTCACTGGCTGGAGGTTATGCGAAAGAATACTTGCG |
| Gene Block 3 | TTCGAAATTCCCGACGATGTGCCGCTGCCAGCCGGTTGGGAGATGGCCAAAACTTCCTCAGGCCAGCGTTACTTCTTAAACCACATAGATCAGACAACTACCTGGCAGGATCCGCGGAAAGCGATGTTAAGTCAAATGAATGTGACGGCACCTACATCCCCGCCAGTCCAGCAGAATATGATGAACTCTGCTAGTGGTCCTCTTCCTGATGGGTGGGAACAGGCGATGACGCAAGACGGCGAAATTTATTACATAAACCACAAGAACAAAACAACCTCGTGGCTTGATCCACGGCTGGACCCACGCTTTGCGATGAATCAG |
| Gene Block 4 | ATGTCTTCAATCCTTCCGTTTACTCCGCCGATCGTCAAGAGGCTATTAGGCTGGAAAAAGGGCGAACAAAATGGGCAGGAAGAAAAGTGGTGTGAAAAAGCCGTCAAATCACTGGTGAAGAAACTGAAAAAAACTGGCCAGCTAGATGAGTTGGAAAAGGCGATTACGACCCAGAATGTAAACACTAAGTGCATCACCATTCCTAGATCTCTGGACGGCCGTCTACAGGTTTCTCACAGAAAGGGGCTACCGCATGTGATCTATTGTAGGCTTTGGCGTTGGCCTGATCTGCATTCCCACCATGAATTAAGGGCTATGGAGCTGTGTGAATTTGCATTCAACATGAAGAAGGATGAGGTGTGCGTTAACCCCTATCATTATCAAAGGGTCGAGACCCCCGTCCTGCCCCCCGTTCTGGTGCCACGTCACACTGAGATACCAGCGGAGTTCCCCCCGCTTGACGACTATTCCCACTCTATCCCAGAGAATACGAATTTTCCGGCTGGTATTGAGCCTCAAAGCAACATCCCAGAGACGCCGCCGCCTGGCTACCTGTCTGAAGATGGAGAGACTAGCGACCACCAGATGAACCATAGTATGGACGCTGGCTCCCCCAATCTAAGCCCCAATCCAATGAGTCCAGCGCACAACAACCTGGATTTGCAGCCGGTCACTTATTGTGAACCAGCATTTTGGTGTTCAATAAGCTACTATGAACTTAACCAAAGGGTTGGAGAGACTTTCCATGCTTCCCAGCCATCCATGACGGTTGATGGATTCACCGATCCCTCTAATTCAGAGAGGTTCTGTTTGGGGCTATTGTCCAACGTTAACCGTAACGCTGCGGTTGAGCTGACTCGTAGGCACATAGGCAGGGGCGTAAGACTTTATTATATT |

| | |
|---|---|
| | GGTGGGGAGGTGTTTGCTGAGTGTCTGTCTGACAGCGCAATTTTC GTCCAGAGCCCGAATTGCAATCAAAGGTATGGATGGCATCCAGC GACGGTCTGCAAGATCCCCCCTGGGTGTAACTTAAAGATCTTTAA TAATCAAGAATTTGCTGCACTTCTTGCTCAAAGTGTGAACCAAGG GTTTGAGGCAGTGTATCAATTAACCAGGATGTGTACGATCCGTAT GAGCTTCGTAAAGGGGTGGGGAGCAGAATACCGTAGGCAGACCG TTACCTCCACACCTTGTTGGATAGAGCTGCACTTAAACGGCCCTCT GCAATGGCTGGATAAAGTTTTAACACAAATGGGTTCACCTTCAAT TAGATGCTCTTCAGTTAGC |
| Gene Block 5 | GGTGGTTCAGCTAGCAGATATAGTCCACCTCCGCCGTACTCTTCTC ATAGTGGATCCGAACAAAAG |
| Gene Block 6 | GGTGGTTCAGCTAGCGAATTGGAATCCCCGCCCCCGCCGTATTCC CGTTACCCTATGGACGGATCCGAACAAAAG |
| Gene Block 7 | GCGCCCGACTCTCCTGATAATGATCTTCGTGCTGGACAGTTCGGC ATAAGTGCCAGGAAGCCATTTACGACCCTGGGCGAAGTCGCGCCC GTATGGGTTCCCGATTCTCAGGCTCCGAACTGTATGAAGTGCGAG GCGAGATTTACGTTTACGAAACGTAGACACCACTGC |
| Gene Block 8 | GAAACTGACCCAGAAGCAGAACCGGAATTAGAGCTATCTGACTA TCCTTGGTTTCACGGGACATTGTCTAGGGTTAAGGCGGCCCAGTT GGTGCTGGCAGGTGGTCCTCGTAACCATGGTTTATTCGTTATCCGT CAAAGTGAAACTAGGCCAGGGGAGTACGTCCTAACTTTTAATTTC CAGGGGAAAGCGAAACACTTGCGTCTTTCTCTAAATGGTCACGGA CAGTGCCATGTCCAACACCTTTGGTTTCAATCTGTTCTTGACATGT TAAGGCACTTTCATACACATCCTATACCTCTGGAGTCCGGTGGTA GCGCCGATATTACACTTCGTAGCTACGTGCGTGCCCAGGACCCTC CTCCCGAACCAGGGCCAACC |
| Gene Block 9 | GGTACCCTATTATAATTCATCGTGTTCAAACCATGGAAACATGGT TTCGAACGCCTGGTGAATCTCAGCGAAGGAGGGTCTATCTGATGG ATTCCACTGCCAGCACGCACGCATTAATTCGTATACTTTTTCGGGA CAGCCCTCAGGACGTTCCATACGATAATCTTTTTCCAGTAGCTCGT ACACTTGAGACAGGTCTATCCCGGGATAAGGAGACATTCCGTAGG |

| | |
|---|---|
| | TGGCAATCTCCCATAGCAGCACTCCAAATGCCCAGACGTCTGACT TAATGCTGAACTTATTATAAGCTAGTGATTCCGGCGCTGTCCATTT AATGGGAAATTTTGCACCAGCATGGGCGGTATAGGTATCTCCAGT CATAAGCCTGGAAAGTCCGAAATCAGCCACCTTGACAAGGTGATT CTCACCAACTAAGCAGTTTCTGGCCGCTAAGTCTCTGTGTATGAA ATTTTTCTTTTCCAGATATTCCATTGCACTGCTTATCTGGGTAGCC ATGTAAAGTAAAACCACGGCATTGACTTCCTGACGATTACACTCT CTAAGATAGTCAAGCAAGTTGCCATATGTCATAAACTCTGTAATA ATATAAAACGGGGGCTCCCTAGTACATACCCCCAACAGCTGAACT AGATTCGGATGTTTAATTTCCTTCATAACAGCCGCTTCTTTCAAGA ACTCTTCAACTTCCATTGTATCTTCTTTCAACGTCTTTACGGCAAC CGTTAGGGAGTACTTCTTCCATACTCCTTCGTATACTTCGCCGTAC TGACCCCCTCCTAGTTTGTGTTTCATTGTAATGTCTGTACGTTCCA TCTCCCAGGCTAGTACGCTCGCGATTACGGAGAAGATACTAAAAC AACGTAGAAGCTGCATCCTAGGAATTCCTTGTTGAATTCGAATTT TCAAAAATTCTTACTTTTTTTTTGGATGGACGCAAAGAAGTTTAAT AATCATATTACATGGCATTACCACCATATACATATCCATATACAT ATCCATATCTAATCTTACTTATATGTTGTGGAAATGTAAAGAGCC CCATTATCTTAGCCTAAAAAAACCTTCTCTTTGGAACTTTCAGTAA TACGCTTAACTGCTCATTGCTATATTGAAGTACGGATTAGAAGCC GCCGAGCGGGTGACAGCCCTCCGAAGGAAGACTCTCCTCCGTGCG TCCTCGTCTTCACCGGT |
| Gene Block 10 | GAATTCATGCAATTGTTGAGATGTTTCTCTATCTTCTCTGTTATCG CTTCTGTTTTGGCTCAGGAACTGACAACTATATGCGAGCAAATCC CCTCACCAACTTTAGAATCGACGCCGTACTCTTTGTCAACGACTA CTATTTTGGCCAACGGGAAGGCAATGCAAGGAGTTTTTGAATATT ACAAATCAGTAACGTTTGTCAGTAATTGCGGTTCTCACCCCTCAA CAACTAGCAAAGGCAGCCCCATAAACACACAGTATGTTTTTTATC CGTACGACGTTCCAGACTACGCTGGTGGTGGTGGTTCCGGTGGTG GTGGTTCTGGTGGTGGTGGTTCAGCTAGCTTCAAGGGTTCTACTG CTGAAAACGCTGAATACTTGAGAGTTGCTCCACAATCTTCTGAAT |

| | |
|---|---|
| | TTGGATCCGAACAAAAGCTTATCTCCGAAGAAGACTTGTTCGAACACGACGAATTGGAGGGAAGGGGATCTTTGCTAACGTGCGGGGATGTCGAAGAAAACCCTGGCCCTATGCAGTTGTTAAGATGCTTCTCTATCTTTTCCGTTATCGCTTCTGTTTTAGCGCAGGAACTTACTACAATTTGTGAACAAATACCTAGTCCCACGTTGGAGTCAACACCGTACTCCCTGTCTACTACAACAATACTTGCGAACGGTAAGGCCATGCAAGGAGTCTTTGAATACTACAAGTCAGTTACCTTCGTATCTAACTGCGGGAGTCACCCGAGCACCACATCCAAGGGGAGTCCTATAAACACTCAGTATGTATTCGACTATAAGGATGACGATGACAAGGGCGGCGGAGGCTCCGGTGGAGGCGGAAGCGGCGGTGGAGGTAGTACTAGTGCGACCGTGAAATTTAAATATAAAGGCGAAGAAAAACAGGTGGATATTAGCAAAATTAGACGGGTGCGTCGCAAGGGCAAATGCATTAGGTTTTACTATGATCTGGGCGGCGGCAAATATGGCAGGGGCATAGTGAGCGAAAAAGATGCGCCGAAAGAACTGCTGCAGATGCTGGAAAAACAGAAAAAAGTCGACTTTGAGCATGATGAGTTGTAATAGCTCGAG |

**Table S4**. List of oligonucleotide primers

| Primer | Sequence |
|---|---|
| Pf1 | GATTACCCTAGGATGCAGACTCTCCTTGTGAGCTCGCTT |
| Pr1 | GAGCACCATATGCTGCATGGAAGCATAATCTTCCAAGATAG |
| Pf2 | TCACATCCTAGGCAACTTACCCCTACCTTCTAC |
| Pr2 | GACCGTCATATGAGAGTTGGAGTTCACCACCC |
| Pf3 | TCGGCACCTAGGATGGTTTTCACCTTAGAGGATTTC |
| Pr3 | TCGTCACATATGCGCAAGTATTCTTTCGCATAACC |
| Pf4 | GAAGTCGCTAGCATGGCGACCGTGAAATTTAAATATAAAG |
| Pr4 | GCACTTGGATCCTTTTTTCTGTTTTTCCAGCATCTGCAG |
| Pf5 | GAAGTCGCTAGCATGGCGACCGTGAAATTAAAATAT |
| Pf6 | TCGGCAGCTAGCATGGTTTTCACCTTAGAGGATTTC |
| Pr6 | TCGTCAGGATCCCGCAAGTATTCTTTCGCATAACC |
| Pf7 | TCCGGTGGTGGTGGTTCTGGTGGTGGTGGTTCAGCTAGCGCAG AGTGGCCATTACGGCC |
| Pr7 | CTCCAAGTCTTCTTCGGAGATAAGCTTTTGTTCGGATCCGAGGC CGAGGCGGCC |
| Pf8 | GGTGGTGGTGGTTCTGGTGGTGGTGGTTCAGCTAGCTTCGAAA TTCCCGACGATGTGC |
| Pr8 | CAAGTCTTCTTCGGAGATAAGCTTTTGTTCGGATCCCTGATTCA TCGCAAAGCGTGG |
| Pf9 | AGCACTGCTAGCATGTCTTCAATCCTTCCGTTTACTC |
| Pr9 | ACGTGCGGATCCGCTAACTGAAGAGCATCTAATTGAA |
| Pf10 | GGTGGTGGTGGTTCTGGTGGTGGTGGTTCAGCTAGCAGA |

| | |
|---|---|
| Pr10 | CAAGTCTTCTTCGGAGATAAGCTTTTGTTCGGATCCACTATG |
| Pf11 | GGTGGTGGTGGTTCTGGTGGTGGTGGTTCAGCTAGCGAA |
| Pr11 | CAAGTCTTCTTCGGAGATAAGCTTTTGTTCGGATCCGTCCAT |
| Pf12 | GTTGGAGCTAGCGCGCCCGACTCTCCTGAT |
| Pr12 | GTTGGAGGATCCGCAGTGGTGTCTACGTTTCG |
| Pf13 | CGTCCGGCTAGCGAAACTGACCCAGAAGCAGAAC |
| Pr13 | GGTCGTGGATCCGGTTGGCCCTGGTTCGGG |
| Pf14 | GGTCGGGCTAGCGGACTCAGATCTAGTGGCAGC |
| Pr14 | GGTCGTGGATCCTGCCCCATAATCGGGTTCAGC |
| Pf15 | GGTTACGCTAGCCGCCTTTCAGAGCCTGTTCCA |
| Pf16 | GCTATCGGTACCCTATTATAATTCATCGTGTTCAAACC |
| Pr16 | GTACTGACCGGTGAAGACGAGGACGCACGGAGG |
| Pf17 | CTGACGGAATTCATGCAATTGTTGAGATGT |
| Pr17 | CTGATCCTCGAGCTATTACAACTCATC |
